# Remodelling of the endothelial cell transcriptional program via paracrine and DNA-binding activities of MPO

**DOI:** 10.1101/2023.05.30.542845

**Authors:** Ruiyuan Zheng, Kyle Moynahan, Theodoros Georgomanolis, Egor Pavlenko, Simon Geissen, Athanasia Mizi, Simon Grimm, Harshal Nemade, Rizwan Rehimi, Jil Bastigkeit, Jan-Wilm Lackmann, Matti Adam, Alvaro Rada-Iglesias, Peter Nuernberg, Anna Klinke, Simon Poepsel, Stephan Baldus, Argyris Papantonis, Yulia Kargapolova

## Abstract

Myeloperoxidase (MPO) is an enzyme that functions in host defence by catalysing the formation of reactive oxygen intermediates. Synthesized majorly by myeloid progenitor cell types and neutrophils, MPO is released into the vascular lumen during inflammation, where it may adhere and subsequently enter endothelial cells coating vascular walls. Here, we show that MPO actually enters the nucleus of these endothelial cells and binds chromatin independently of its enzymatic activity to cause changes in chromatin structure. At its binding sites, MPO drives chromatin decondensation, while enhancing condensation at flanking regions. We further show that MPO binds loci relevant for the activation of the endothelial-to-mesenchymal transition (EndMT) and the migratory potential of ECs. Finally, MPO interacts with the RNA-binding factor ILF3 affecting its relative abundance between cytoplasm and nucleus. This leads to ILF3:MPO- driven transcriptional and post-transcriptional regulation. Accordingly, MPO-knockout mice show reduced EC numbers at scars formed after myocardial infarction, indicating reduced neovascularization. In summary, we describe a non-enzymatic role for MPO in coordinating EndMT and controlling the fate of endothelial cells through direct chromatin binding and association with such co-factors as ILF3.

## INTRODUCTION

Myeloperoxidase (MPO), a lysosomal protein, functions in response to pathogens during bacterial and fungal infections and contributes to pathogen clearance within the phagosome. Catalytically active MPO uses hydrogen peroxide and halide or pseudohalide ions to catalyse production of acids that are highly reactive and cause oxidation of proteins and small molecules, (Davies & Hawkins, 2020; Davies, 2011). Multiple studies associated products of enzymatic activity of MPO with cardiovascular diseases, multiple sclerosis, cancer and atherosclerosis (Kargapolova *et al*, 2021a).

Apart from its microbicidal function, MPO has been shown to modulate endothelial cell (EC) surface integrity and EC function by the reduction of NO-bioavailability (Baldus *et al*, 2001). It also acts as an autocrine regulator of leukocyte function by modulating leukocyte activation and adhesion (Nussbaum *et al*, 2013). Moreover, it delays neutrophil apoptosis *in vitro* (El Kebir *et al*, 2008), suggesting that it functions as a survival signal for neutrophils and as such may prolong the duration of an inflammatory response.

Along with the well-studied enzymatic activity of the protein, MPO possesses non-enzymatic properties. First of them is an intrinsic affinity to DNA. In the process of formation of neutrophil extracellular traps (NETosis), MPO, which is in normal conditions primarily found in granules of resting cells, translocates to the nucleus where it causes the citrullination of histones and chromatin decondensation (Metzler *et al*, 2011; Pilsczek *et al*, 2010; Papayannopoulos *et al*, 2010). Second example of non-enzymatic activity of MPO has been demonstrated by Manchanda and colleagues (Manchanda *et al*, 2018) and was attributed to the cationic charge of the protein, causing a collapse of the endothelial glycocalyx.

Apart from its activity on the endothelial cell surface, MPO has a capacity to enter endothelial (EC) and epithelial cells (Baldus *et al*, 2001; Astern *et al*, 2007). In less than six hours, MPO can be detected in the cytoplasm and the nucleus of ECs, presumably remaining enzymatically active (Yang *et al*, 2001). However, ∼40% of the MPO is inactive at the site of inflammation (Bradley *et al*, 1982). For instance, studies have shown the presence of 16–29 mg/ml of enzymatically inactive MPO (iMPO) in arthritic joints (Edwards *et al*, 1988). Interestingly, both, inactive and active MPO can drive transcriptional changes in ECs (Lefkowitz *et al*, 2000). Among others, the expression and secretion of cytokines interleukin (IL)-6 and -8, MCP-1, and GM-CSF increases rapidly and in a dose-/time-dependent manner following MPO treatment (Lefkowitz *et al*, 2000). These findings suggest that MPO might affect gene expression via activity-independent mechanisms. Provided an affinity for chromatin, it may act as a transcriptional activator or repressor via direct regulation of chromatin organization. In spite of accumulating evidence supporting its non-enzymatic function on gene expression, the exact mechanism of how MPO acts, remains unknown. Here, we sought to shed new light on the nuclear roles of MPO, likely constituting the first example of an enzyme contributing to cell-to-cell communication by entering the nucleus and directly binding chromatin to regulate gene expression.

## RESULTS

### MPO accumulates in endothelial cell nuclei within hours

To assess MPO transmigration into EC nuclei, we treated human umbilical vein ECs with MPO for 2 and 8 hours and performed immunofluorescence and fractionation western blot experiments. Accumulation of MPO in the nuclei of ECs 2 hours after treatment was microscopically apparent (Figure 1A,B). This was reflected in the respective quantified mean nuclear signal intensities measured in >100 cells (Figure 1A, lower panel). At 8 hours post-treatment, nuclear MPO levels were ∼40% less than those at 2 hours (Figure 1B). Notably, nuclear accumulation of MPO resulted in its presence on the chromatin fraction already at 2 hour post-treatment (Figure 1C). Catalytically-inactive MPO was similarly enriched in the chromatin fraction of HUVECs (Figure 1D,E), indicating that this occurs independently of catalytic activity.

**Figure 1.**
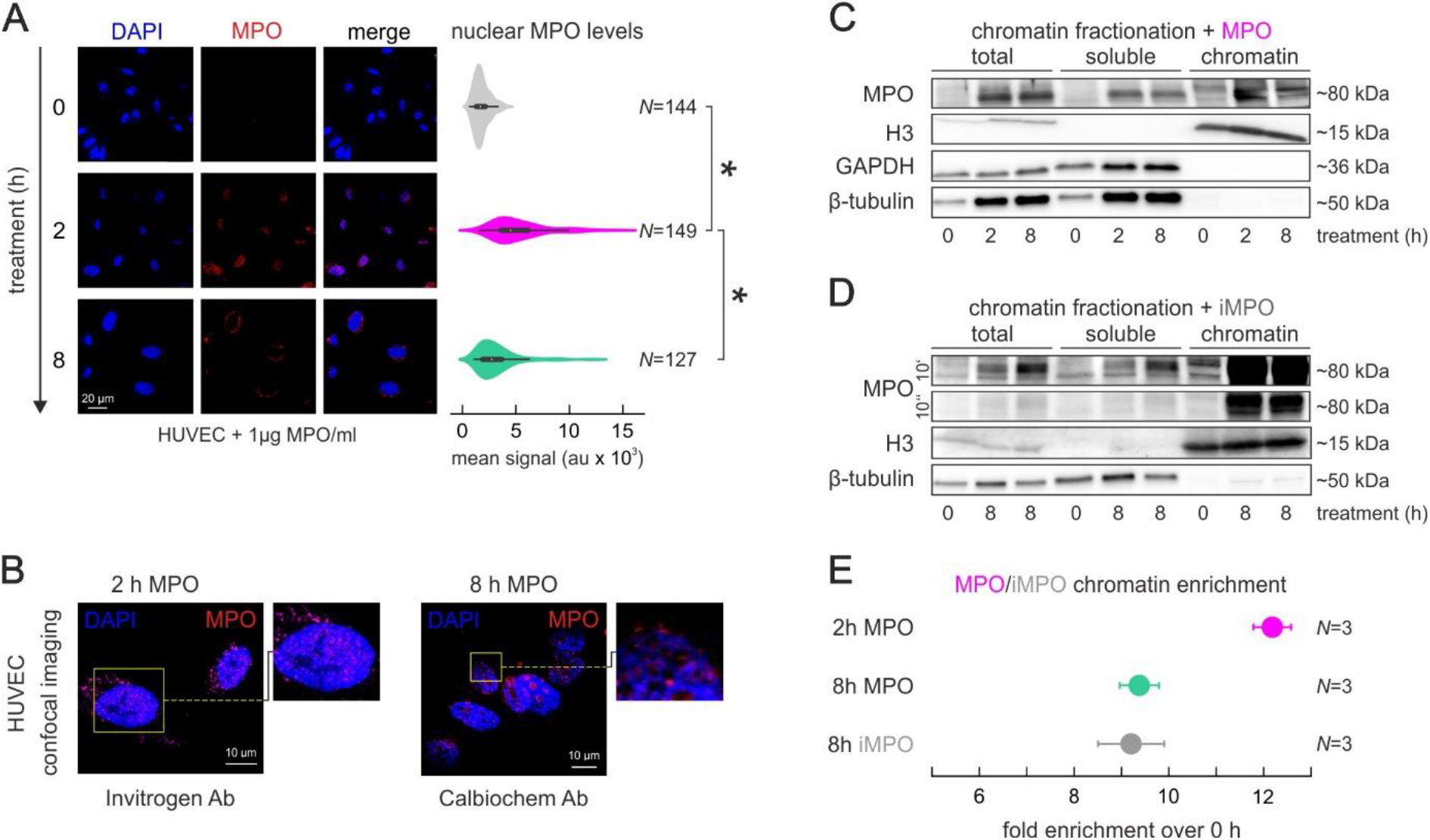
MPO translocates into ECs and binds chromatin already within 2 h after treatment. (A) Representative images with immunofluorescence signal of MPO in ECs after 0, 2 and 8 hours of MPO added to the media, quantification of mean nuclear MPO signal (right panel) calculated for 10-15 random images of three biological replicates in form of violin plots, *N* – number of individual nuclei; scale bar = 20μm. (B) Confocal image of MPO translocation in ECs upon 2- and 8-h MPO treatment using two different antibodies for IFs in at least 3 biological replicates; scale bar = 10μm. (C) Western blot determining total, soluble, and chromatin-bound MPO levels in ECs treated 2 or 8 h. (D) Western blot determining total, soluble and chromatin-bound levels of MPO in ECs treated 8 h with iMPO; images taken for 10 sec or 10 min. (E) Quantification of chromatin-bound MPO and iMPO enrichment compared to untreated cells and normalized to histone H3 levels; *N* = 3 biological replicates.

We sought to exclude potential effects on ECs caused by the MPO enzymatic activity through activation of autophagy by accumulation in lysosomes (Yogalingam *et al*, 2017) or of signalling pathways on the cell surface during its translocation into the nucleus (Khalil *et al*, 2018). Western blot analysis of key autophagy factors revealed no difference in their levels (i.e., ATG5, ATG12, ATG7 or LC3A/B in Figure S1A). Consistently, no TFEB translocation into the nucleus could be detected by immunofluorescence (Figure S1D). Furthermore, levels of latent monomeric TGFβ remained unaltered upon MPO treatment, and expectedly increased in hydrogen peroxide-treated ECs (Figure S1A,C). A mild decrease of phospho-SMAD, rescued by the addition of catalase, and no alteration in phospho-ERK levels were observed after MPO treatment (Figure S1B,C). This indicates that MPO-driven effects on transcription regulation of endothelial cells likely happen independently of signalling pathways, affected by MPO treatment.

### MPO directly binds chromatin at nucleosome-free regions

To estimate the propensity of MPO to biind nucleosomes *in vitro* electromobility shift assays (EMSA) were performed, in which MPO was titrated between 50 and 1000 nM to bind nucleosomes with ‘no’ (0bp) and ‘long’ (∼100bp) DNA linkers (Figures 2A and S2A). MPO displayed binding to long linker nucleosomes at low titers, and ‘no linker’ nucleosomes at medium to high titers (Figure 2A, lower panel). Importantly, enzymatic activity was not required for efficient binding (Figure S2B).

**Figure 2.**
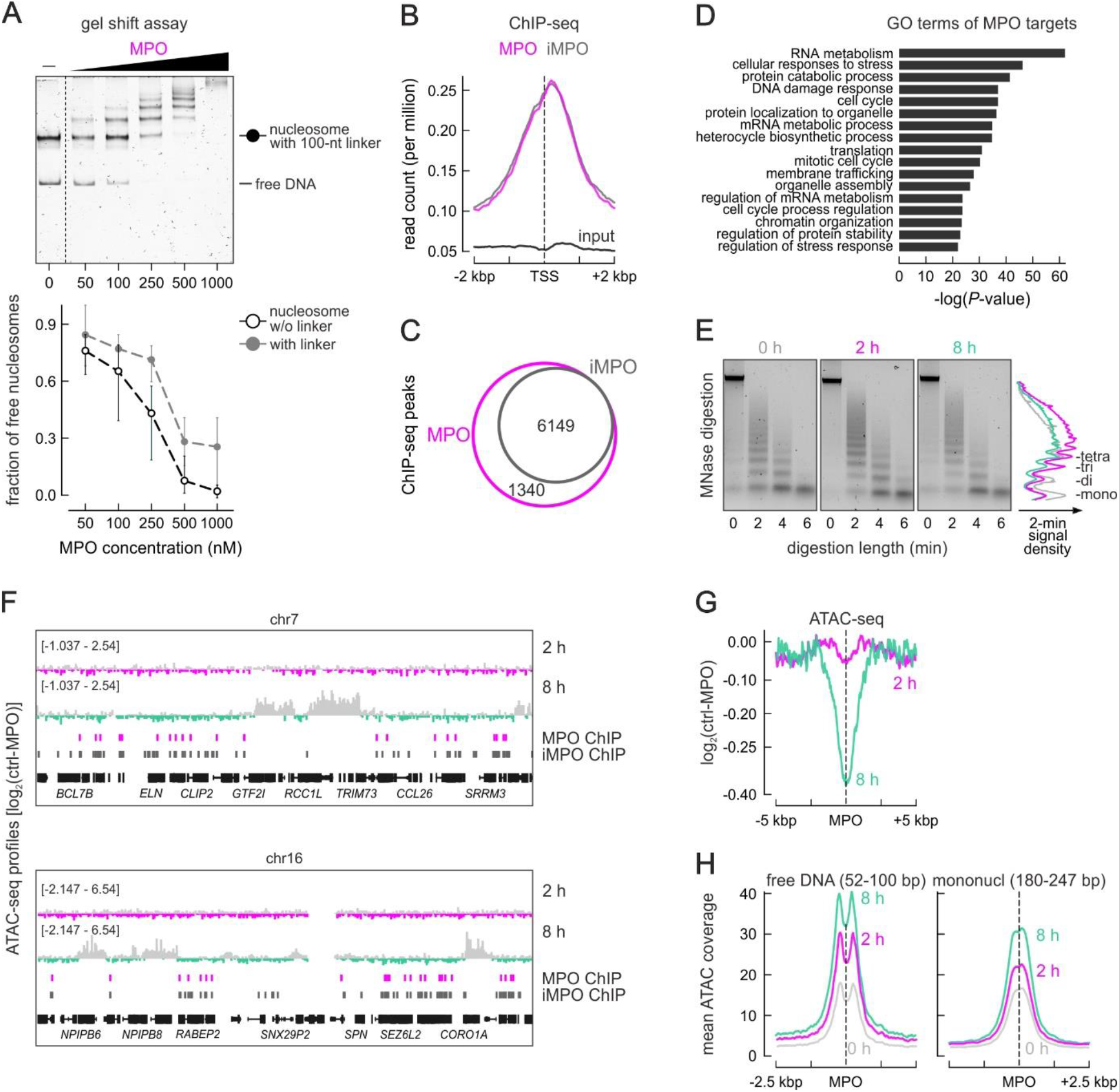
Myeloperoxidase binds to chromatin in vitro and in cyto, causing changes in chromatin condensation. (A) EMSAs performed with increasing titers of MPO (50-1000nM) and nucleosomes with 100bp, long liker DNA or with 0bp, ‘no linker’ DNA. Quantification of the assay is shown in the lower panel, 3 replicas were analysed. (B) Line plot showing mean distribution of MPO (pink) and iMPO (grey) ChIP-seq and input signal (black). (C) Venn diagrams showing overlap between conserved ChIP-seq peaks for MPO (pink) and iMPO (grey). (D) Significantly enriched GO terms associated with sets of 3000 genes, bound by MPO (within 670 bp of TSSs). (E) Electrophoresis profiles of DNA fragments released by MNase from ECs treated with MPO for 0, 2 and 8 h. Signal density after 2 min of MNase treatment was quantified and plotted as a line-plot (right panel) – mono-, di-, tri, and tetra-nucleosomes are labeled; experiment performed in triplicates. (F) Representative genome browser views of ATAC-seq coverage-based log2 ratios of control vs 2-h or control vs 8-h MPO-treated chromatin merged with positions of MPO and iMPO ChIP-seq peaks (G) ATAC accessibility in the 10 kbp around MPO-bound positions calculated as log2 ratios of control vs 2-h MPO (pink) or control vs 8-h MPO treatment (green). (H) ATAC accessibility in the 5 kbp around MPO-bound positions for a subset of reads between 52 and 100 bp, corresponding to free DNA (left) or between 180 and 247 bp, corresponding to mononucleosomes (right).

The genome-wide capacity of chromatin binding by enzymatically-active and inactive MPO (iMPO) was assessed by chromatin immunoprecipitation and sequencing (ChIP-Seq; Figure 2B). To account for the transient nature of MPO binding to chromatin, this experiment was performed at 8 h post-treatment using an antibody successfully applied in immunostainings and western blots. MPO and iMPO binding was mainly detected at transcription start sites (TSSs) (Figures 2B and S2C,E). MPO/iMPO ChIP-Seq was performed in duplicates, with conserved peaks converging for active and inactive protein (Figure 2C and Table S1). For example, MPO/iMPO peaks co-localized with regions of open chromatin at the *KIT* locus (Figure S2D) and were overall enriched for POLR2A and H3K4me3 signal based on the analysis of ENCODE ChIP-Seq data (Figure S2F). This suggests a preference for TSSs of actively transcribed genes, but not for enhancer regions (Figure S2E,G). Gene ontology analysis performed via Metascape (Tripathi *et al*, 2015) uncovered gene clusters related to cell cycle, chromatin organization, and VEGFA-VEGFR2 signaling (Figure 2D), all previously implicated in the process of angiogenesis (Abhinand *et al*, 2016).

The consequences of MPO binding to chromatin were assessed via micrococcal nuclease digestion followed by electrophoretic analysis at different time points after treatment. These experiments indicated increased, time- and concentration-dependent chromatin condensation caused by MPO treatment (Figures 2E and S2H). To map sites of MPO-driven chromatin condensation, ATAC-seq was performed. Little change in chromatin accessibility (with a bias for increased condensation) was measured 2 h post-treatment and globally distributed, as exemplified for chromosomes 7 and 16 (Figure 2F). Longer MPO treatment caused profound condensation at specific regions and moderate de-condensation next to MPO binding sites (Figure 2F,G). The specific areas of moderately increased chromatin condensation were computationally identified as areas of 2-fold condensation in a 2-kbp or larger windows (Table S2). According to the nucleosome positioning analysis, MPO-bound regions are mostly nucleosome-free of with binding close to the entry/exit site of the adjacent nucleosome. However, MPO binding did not cause any shift in nucleosome positioning (Figure 2H).

### MPO induces Endothelial-to-Mesenchymal transition to endothelial cells

The impact of MPO treatment on gene expression was measured by RNA-seq at 2 and 8 hour post-treatment compared to untreated ECs. Differential expression of 456 and 65 genes (fold change > 2; Padj < 0.05) was observed after 2 and 8 h of MPO treatment, respectively (Figure 3A), suggesting maximum degree of gene regulation coincides with maximum of chromatin-bound MPO (Table S3). GO term enrichment revealed ossification, blood vessel development and negative regulation of Notch signaling for the upregulated genes, and MAPK family signaling as well as chloride transmembrane transport for downregulated ones at 2 hours after treatment (Figure 3A). Gene Set Enrichment Analysis (GSEA) revealed β-catenin signaling, adipogenesis, DNA repair and other pathways were shared by genes regulated at both 2- and 8-h post-treatment (Figure S3B,C). Treatment of ECs with enzymatically inactive mutant variant of MPO, Q91T (Moguilevsky et al, 1991) allowed identification of only 42 differentially-expressed genes (fold change > 2; Padj < 0.05) of which 31 were also significantly regulated by enzymatically-active MPO treatment (Table S3). Genes related to the Senescence Associated Secretory Phenotype (SASP) were regulated by both the active and the mutant form of MPO (Figure S3B).

**Figure 3.**
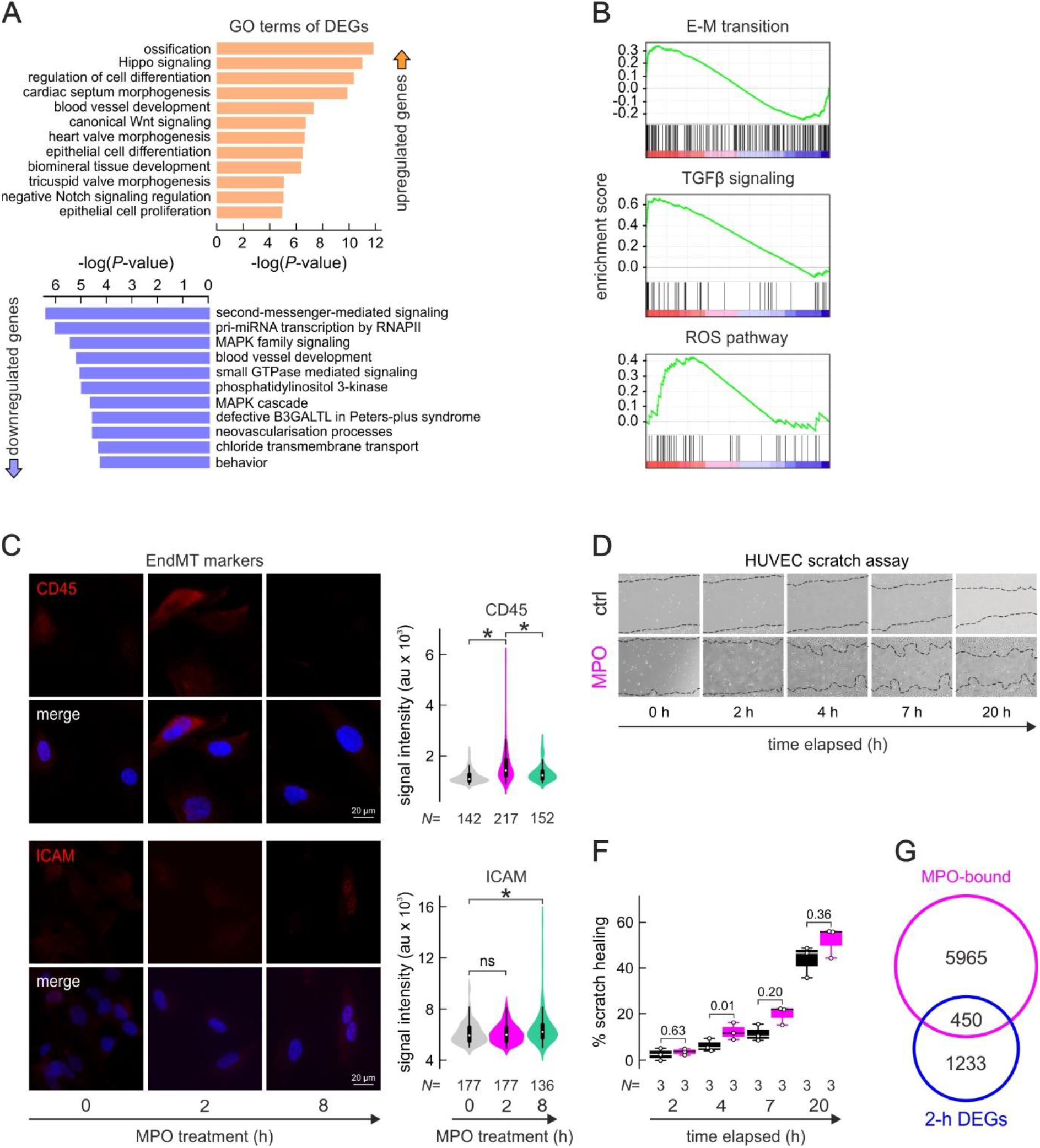
MPO triggers endothelial-to-mesenchymal transition in ECs. (A) GO terms associated with up- (orange) or down-regulated (blue) genes upon 2-h MPO treatment. (B) Gene set enrichment analysis (GSEA) of ranked gene expression data, generated by RNA-seq of HUVECS under 2 h MPO treatment, FDR q-values < 0.1. (C) Representative immunofluorescence images of CD45 and ICAM in HUVECs upon 0, 2 and 8 h of MPO treatment, scale bar=10μm. Mean signal of total ICAM and CD45 plotted as violin plots – results of three biological replicates are summarized, N – number of individual cells. (D-F) Wound healing assay performed after MPO (top row) or mock treatment (bottom row) over the period of 20 hours, images taken at 0, 2, 4, 7 and 20 h, % of scratch healing calculated relatively to 0 time point, mean data of three biological replicates are plotted. (G) Venn diagrams showing overlap between MPO-bound genes and differentially expressed genes at 2 h post treatment.

When up- and down-regulated genes were co-considered, GO analysis suggested that ECs undergo endothelial-to-mesenchymal transition (EndMT). Selected markers of endothelial and mesenchymal cells were assessed by RT-qPCR (Figure S3A) and immunofluorescence (Figure S3D), and confirmed EndMT induction independent of enzymatic activity of MPO. Protein levels of CD45, an indispensable driver of the EndMT following myocardial infarction (Bischoff *et al*, 2016; Kovacic *et al*, 2019), increased 2 and 8 h post-treatment in ECs (Figure 3C,D). CD34, an endothelial progenitor marker, was significantly downregulated 2h post-treatment (Figure S3D,E), whereas CD31 levels (platelet/endothelial cell adhesion molecule-1 - PECAM- 1) increased at 8 h post-treatment (Figure S3D,E). This shows that MPO triggers partial and transient EndMT to endothelial cells.

A hallmark of EndMT is increased migratory potential. The wound healing/scratch assay, performed with and without MPO in the media, showed MPO treatment enhanced migratory potential of ECs and supported hypothesis that cells acquired mesenchymal-like properties (Figure 3E,F). In comparison to other factors involved in EndMT, e.g. tumor necrosis factor alpha (TNFα) a proinflammmatory cytokine that signals through nuclear factor kappa B (NF-κB) (Diermeier *et al*, 2014), MPO-driven transcriptional signatures were distinct, with no significant overlap (Figure S3F) and with just 32 genes being regulated by both treatments (2 fold change, Pval < 0.05), of which 9 were reciprocally regulated (Table S3). Further, we compared TSSs bound by MPO (within <700 bp) and identified 450 genes that were transcriptionally regulated by at least 50%. Similarly, genomic intervals identified to undergo chromatin condensation were crossed with the gene set regulated at 8 h post-treatment and 136 sites were annotated inside or flanking differentially regulated genes (Padj < 0.05). Both findings suggest that MPO can directly affect gene expression via the induction of changes in chromatin structure, mostly around gene promoters.

### MPO interacts with ILF3 and affects its subcellular localization

To determine interaction partners of MPO in EC nuclei, an immunoprecipitation of MPO from nuclei treated for 0 and 8 h was performed (Figure 4A). The MPO interactome was catalogued by mass spectrometry and visualized using Metascape. MPO interacts with proteins known to regulate mRNA processing and stability, nucleosome positioning, and pre-rRNA complexes (Figure 4B). One of the RNA-binding factors identified amongst interacting partners of MPO (and confirmed by co-IP western blot) was ILF3 (Figure 4A,D). An analysis of motifs at MPO-binding sites revealed ILF3- and PCBP1-like motifs among the most plausible (Figure 4C), and both these proteins were identified as MPO-interacting partners (Table S4).

**Figure 4.**
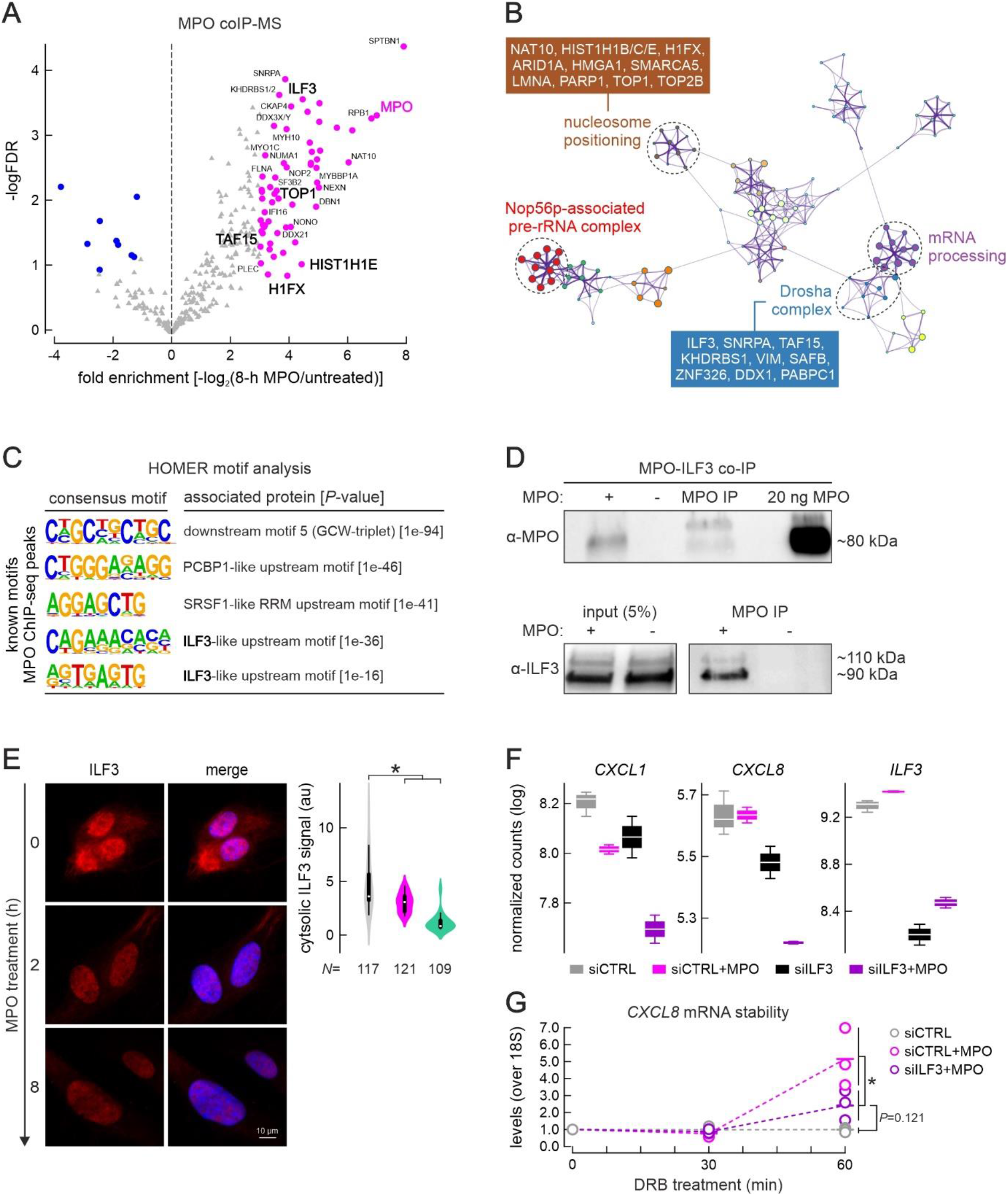
MPO binds to transcription factor ILF3 and regulates specific gene stability and expression. (A) Volcano plot of MPO interacting proteins, significantly enriched factors are depicted in pink. (B) Metascape-based network representation of GO terms associated with proteins co-purifying with MPO, exemplified are proteins forming GO terms “Large Drosha complex” and “nucleosome positioning”. (C) Five most relevant motifs found with help of HOMER and within a 200 bp region around the center of MPO ChIP-Seq peaks, resembling motifs recognized by PCBP1, SRSF1 and ILF3 proteins. (D) Western blot visualization of ILF3 protein co-immunoprecipitated with MPO. (E) Immunofluorescence image depicting decrease of ILF3 signal in the cytoplasm of MPO-treated ECs, with quantification (right panel), scale bar=10μm. (F) Box plots, showing representative regularized log transformed counts of 3’-end sequencing experiment of Cxcl1, Cxcl8 and Vegfa upon treatment with mock control siRNA, siRNA against ILF3, or a combination of siRNA treatments with MPO. (G) Stability assay performed with help of RT-qPCR and after inhibition of transcription with 5,6- Dichlorobenzimidazole 1-beta-D-ribofuranoside (DRB) for 1 hour time, detecting levels of Cxcl8 transcript, and normalized by 18S.

ILF3 is a well-characterized RNA-binding protein, with its smaller isoform (NF90) recently shown to also act as a transcription factor (Reichman *et al*, 2003; Wu *et al*, 2018). ILF3 localization via immunofluorescence showed loss of cytoplasmic ILF3 signal at 2 and 8 h post-treatment (Figure 4E). Additionally, gain of NF90 in the chromatin fraction of ECs at 2 and 8 h post-treatment was detected (Figure S4A). Interestingly, a slight increase of NF90 in the chromatin fractions of ECS was also observed after treatment with the enzymatically inactive MPO variant Q91T (Figure S4A), suggesting that changes in ILF3 abundance were not linked to the enzymatic activity of the protein.

To determine the transcriptional network controlled by ILF3, we performed ILF3 knockdowns (with an efficacy of >70%; Figure S4B) followed by 3’-end sequencing of RNAs. ILF3 depletion caused significant changes to the levels of 110 genes (Padj < 0.1; Table S5 and Figure S4C). Interestingly, when MPO treatment was performed on top of ILF3 depletion, only 47 genes were differentially expressed (Table S5). Then, MPO treatment led to the differential expression of 37 genes when performed upon control siRNA transfection, whereas only 13 genes were significantly regulated by MPO after ILF3 knockdown, suggesting that partial regulation of genes via MPO is dependent on sufficient ILF3 levels.

Finally, we speculated that transcripts known for being stabilized by ILF3 (Vrakas *et al*, 2019), e.g. *CXCL8* (Figure 4F) and *PPDPF* (Figure S4D), is downregulated by ILF3 depletion, but MPO treatment led to their stabilization. Thus, we measured transcript abundance after inhibition of ongoing transcription by DRB (5,6-Dichloro-1-β-D-ribofuranosylbenzimidazole). Monitoring the effects of DRB inhibition on *CXCL8* and *PPDPF* abundance for up to 60 minutes revealed that MPO treatment increased stability of both, whereas the combination of MPO treatment with ILF3 depletion significantly diminished this effect (Figure 4G and S4E). These effects exemplify one possible mode-of-action of MPO for gene expression remodelling of ECs.

### *In vivo* role of MPO in neovascularization in a model of myocardial infarction

Endothelial cell migration is essential for angiogenesis, a formation of new vessels from pre-existing ones. This process is regulated by a tight balance between pro- and antiangiogenic agents and involves a cascade events, of which migration of capillary endothelial cells is an essential component. A recent publication by Tombor and colleagues revealed that after myocardial infarction (MI) ECs acquire a transient state of mesenchymal transition in the first week after MI (Tombor *et al*, 2021). Interestingly, inhibition of mesenchymal activation by the TGFβ inhibitor Galunisertib (LY2157299) reduced the incidence of clonal expansion, suggesting that mesenchymal activation contribute to endothelial expansion (Tombor *et al*, 2021). Curiously, as a result of Galunisertib treatment, a reduced serum levels of MPO, IL-1β and IL-6 were observed in the acute pancreatitis model, suggesting those factors could also contribute to mesenchymal activation (Liu *et al*, 2016). We overlapped TGFβ regulated genes (Glaser *et al*, 2020) with those changes by MPO treatment, and observed activation of expression of 65 common genes and repression of 71 common genes by MPO and TGFβ in ECs, whereas 12 genes were reciprocally regulated (Figure 5A). Intriguingly, the cohort of common upregulated genes was enriched in category ‘blood vessel development’ (Figure 5B). Similarly to MPO, TGFβ treatment of ECs suppresses *Cxcl1* and activates and *Icam1* expression, level of *Ilf3* however slightly decreases upon TGF-ß treatment (Figure S3A and S5A). Interestingly, genes bound by MPO directly and positively regulated by MPO and TGF-ß are enriched in ‘Integrin 3 pathway’, ‘signalling by TGF- ß family members’ and ‘EMT’, suggesting MPO can enhance activation of TGFβ signaling by direct chromatin-binding and gene activation (Figure S5B).

**Figure 5.**
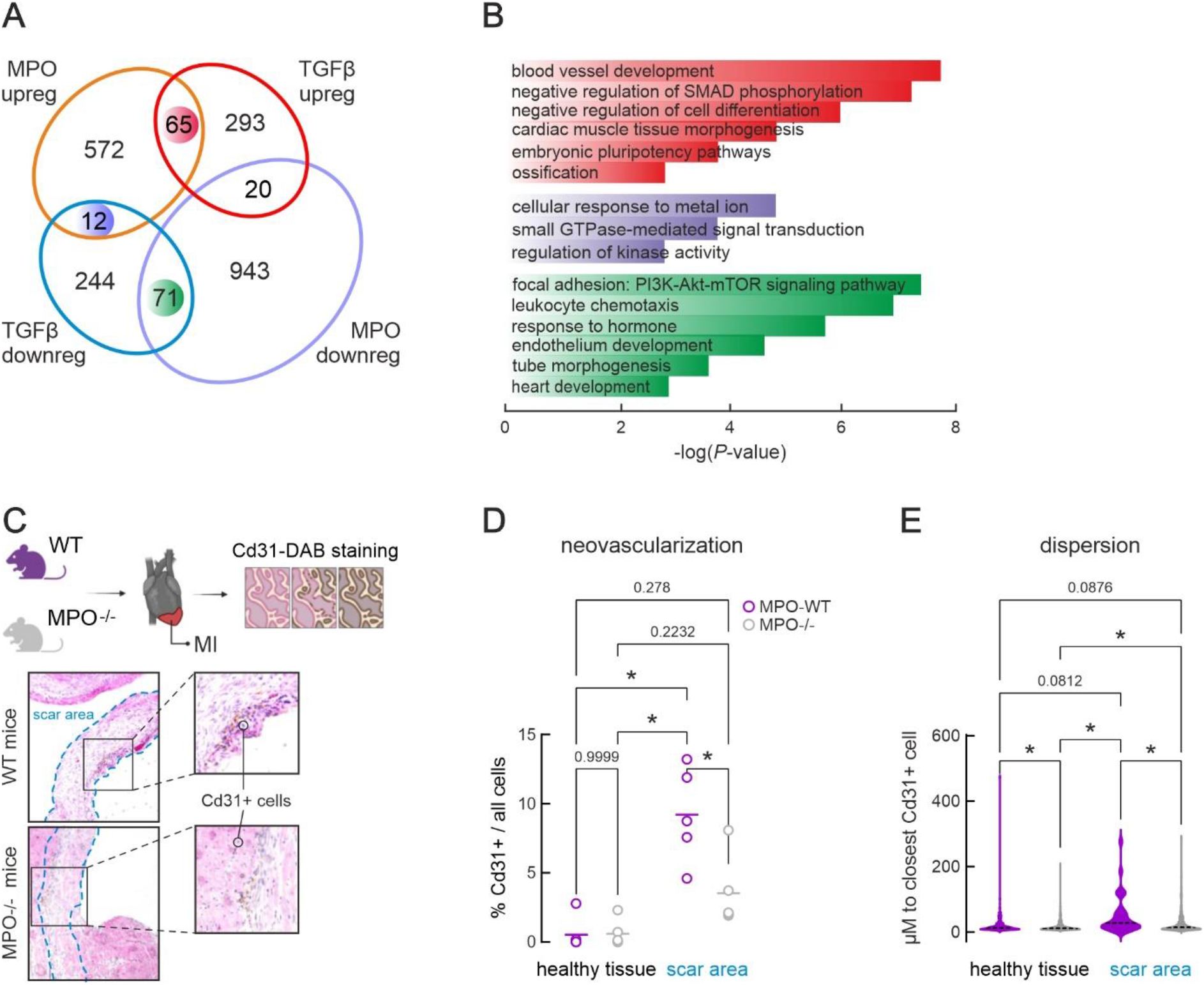
MPO and TGFβ regulate similar sets of genes involved in blood vessel development. (A) Venn diagram depicting the overlap between gene sets, up- and down-regulated upon MPO (current study) and TGFβ treatment (Glaser et al, 2020). (B) Barplot depicting top results of Metascape analysis of gene sets reciprocally or commonly up- or down- regulated by TGFβ (Glaser et al, 2020) and MPO (this study). (C) DAB and H&E staining applied to identify CD31+ cells. (D) Barp lot showing the percent of CD31+ (a measure of capillary endothelial cells) as approximation for neovascularization in healthy and scar tissue of MPO-/- and MPO+/+ animals after 8 weeks post myocardial infarction (2-way Anova test), *: P < 0.001. Each dot represents one animal, >50 images analysed per animal. (E) Bar plot showing distance to the closest CD31+ in healthy and scar regions of MPO^+/+^ and MPO^-/-^ mice after MI, *: P < 0.001.

To ask whether MPO similarly to TGFβ impacts neovascularization, we performed *in vivo* experiment, in which number of CD31^+^ cells, marking capillary ECs, were compared in MPO^-/-^ and MPO^+/+^ mice after 8 weeks post myocardial infarction (Figure 5C). The number of CD31^+^ cells was significantly higher in MPO^+/+^ animals at the scar area, whereas no difference observed for healthy tissue (Figure 5D). Furthermore, distances to the closest CD31^+^ cell was significantly larger in both healthy tissue and scar area of MPO^-/-^ animals (Figure 5E). This observation confirms that MPO might play additional roles in activating EndMT and enhancing cell-to-cell communication coordinated by factors such as TGFβ.

## DISCUSSION

In frame of this work, we have shown that MPO is being endocytosed and transmigrates through the HUVEC cells within a period of 24 hours. By the time of 2 hours post-treatment, MPO is majorly found bound to the chromatin of ECs, and this binding is independent of the enzymatic activity of the protein (Figure 1A-D). *In vitro* and *in cyto* experiments performed in frame of this study suggest MPO preferably binds to the nucleosome-free regions of the chromatin, and at transcription start sites of actively transcribed genes (Figureσ 2A,B and S2C). Locally, at the sites of MPO binding, the chromatin becomes less condensed (Figure 2G). This local de-condensation may change chromatin accessibility for specific transcription factors, thereby affecting transcription of more than four hundred genes (Figure 3G). Interestingly, we have identified increased phosphorylation of histone H1.5 at the position 18 (pS18-H1.5) directly bound by MPO (Supplementary Table 4). The regulation of phosphorylation of Histone 1 variants was previously associated with differentiation of pluripotent cells (Liao & Mizzen, 2017) and is implemented in chromatin de- condensation. On the other hand, regions flanking the MPO binding sites gain condensation (Figure 2F), transcription at those sites gets mainly repressed (Table S2).

During MPO treatment a wave of transcription regulation occurs, peaking at 2 h, which coincides with the maximum of MPO bound to the chromatin. The transcriptional program triggered by MPO leads to partial endothelial-to-mesenchymal transition of ECs and is independent of enzymatic activity of the protein (Figure 3A-F and S3A-E). During this process, alterations in cell morphology, and a gain of a migratory potential are observed (Figure 3E-F). EndMT is a type of cellular transdifferentiation, demonstrated by endothelial cells with remarkable phenotypic plasticity. EndMT partakes in embryonic development and pathogenesis. During the disease of pulmonary arterial hypertension, EndMT is causative for structural and functional pulmonary vessel alterations, during atherosclerosis it leads to an increased vascular wall fibrosis, resulting in vessel narrowing and during fibrodysplasia ossificans progressiva – leads to soft tissue calcification (Piera-Velazquez & Jimenez, 2019). In our setup, the endothelial cells undergo only partial EndMT. It is not driven by altered TGFβ levels (Figure S1C), and is independent of NF-κΒ signaling (Figure S3F). Although TGFβ levels are not altered by MPO, there are genes regulated by both factors (Figure 5A). In particularly, gene set positively regulated by both factors is enriched in ‘blood vessel development’ (Figure 5B). Both factors effect *ILF3* levels, TGFβ represses its expression (Figure S5A), whereas MPO directly interacts with ILF3 protein (Figure 4D) and causes its depletion in the cytoplasm (Figure 4E). Similar to TGFβ, MPO seemingly partakes in mesenchymal activation of endothelial cells *in vivo*, and when lacking, leads to reduced neovascularization after myocardial infarction (Figure 5D).

We describe a non-enzymatic role of MPO in the process of endothelial-to-mesenchymal transition, which we attribute to its activity as a ‘mediator’ protein entering the nucleus and binding chromatin to regulate gene expression. Although, many factors triggering EndMT have been described already, we propose, MPO can be added to this list, and therefore its depletion may beneficially delay the process of EndMT in fibrotic and vascular diseases. Accordingly, MPO deficiency in humans with cardiovascular disease was beneficial for the disease progression and outcome (Teng *et al*, 2017; Rudolph *et al*, 2012).

We observed that MPO drives regulation of SASP transcripts (Figure S3B). This type of regulation might be important during tissue regeneration. The potential role of SASP and senescent cells during wound healing was proposed by Lopes-Paciencia and colleagues (Lopes-Paciencia *et al*, 2019), stating that transient exposure to SASP leads to increased regenerative capacity *in vivo*. Therefore, we speculate that the short exposure of endothelial cells to MPO might be beneficial during the process of tissue regeneration.

Apart from its function on chromatin condensation and directly at TSS in ECs, MPO also binds to a multi-faceted protein ILF3 and causes change in abundance of ILF3 isoforms in the cytoplasm and in the nucleus (Figure 4E, S4A). ILF3, initially well described as an RNA-binding protein, has been implicated in a selective regulation of gene expression, mRNA stabilization of *Bcl2*, *IL2*, *MyoD*, *Cxcl1*, *Il8*, *Vegf* (Vrakas *et al*, 2019), translation inhibition of SASP factors (Tominaga-Yamanaka *et al*, 2012), modulation of viral replication and noncoding RNA biogenesis (Castella *et al*, 2015). Both isoforms of ILF3 (NF110 and NF90) were found in nucleus and cytosol and are shuttling between these two compartments, driven by post-translational modifications (Castella *et al*, 2015). Moreover, (Reichman *et al*, 2003) discovered that ILF3 proteins were associated with chromatin. Wu with co-authors recently performed chromatin immunoprecipitation of NF90/ILF3 from K562 erythroleukemia cells (Wu *et al*, 2018), peaks majorly co-localized with chromatin marks associated with active promoters and strong enhancers. High expression of ILF3 was associated with angiogenesis in cultured human coronary artery endothelial cells (hCAECs), whereas knockdown in hCAECs reduced abundance of several angiogenic factors, and reduced proliferation rate of cells (Vrakas *et al*, 2019). We showed that observed effects of MPO on transcript abundance was dependent on high levels of ILF3, and diminished with ILF3 depletion. We speculate those effects were maintained via ILF3-regulated stability of given transcripts, when combined with MPO, it enhanced post-transcriptional regulation observed.

In conclusion, three modes of enzymatically independent activity of MPO on EC gene regulation were revealed by current work. These include direct effect on transcription, indirect regulation of transcription via events of chromatin condensation/decondensation and lastly, ILF3-dependent regulation of transcripts’ stability.

## MATERIAL AND METHODS

### ChIP-seq and data analysis

HUVEC cells were pre-treated with 1 µg/ml of MPO for 2 or 8 hours; for control experiments fresh media was given to the cells. For each batch of ChIP experiments, ∼12 million cells were crosslinked in 2% PFA for 45 min at 4 °C. From this point onward, cells were processed via the ChIP-IT High Sensitivity kit (Active motif) as per the manufacturer’s instructions, but using the NEXSON protocol for nuclei isolation (Arrigoni *et al*, 2015; Kargapolova *et al*, 2021b). Chromatin was sheared to 200–500-bp fragments on a Bioruptor Plus (Diagenode; 2× 20–26 cycles of 30 s “on” and 30 s “off” at the highest power setting), and immunoprecipitations were carried out by adding 4 μg of the appropriate antibodies (MPO, Calbiochem) to

∼30 μg of chromatin and incubating on a rotator overnight at 4 °C in the presence of protease inhibitors. Following addition of protein-A/G agarose beads and washing, DNA was purified using the ChIP DNA Clean & Concentrator kit (Zymo Research) and used in qPCR or sequencing on a HiSeq4000 platform (Illumina). Where ChIP-seq was performed, at least 20 million reads were obtained, also for the relevant “input” samples. Raw sequencing reads (typically 50 nt-long) were analyzed using the HiChIP pipeline (Yan *et al*, 2014), and peaks were called using MACS2 (Zhang *et al*, 2008). Thresholded MPO ChIP-seq peaks (q-value <0.05) per each cell type and genotype are listed in Supplementary Table 1. For plotting ChIP-seq signal coverage over select regions, ngs.plot was used (Shen *et al*, 2014). The motif search around the MPO binding sites was performed as such: 1) windows of 200 bp of length were defined upstream and downstream of a peak center 2) motif analysis was performed for upstream and downstream selected regions separately, using HOMER software (Heinz et al., 2010) by running findMotifsGenome.pl command; results of de-novo motif discovery were used.

### HUVEC culture and MPO treatment, MPO inactivation

Human Umbilical Vein Endothelial Cells, pooled from several donors, were purchased from Lonza and cultured in FBS reduced Endopan 3 media. HUVEC cells of passages 5-10 were used for experiments, when cells were actively proliferating. Culturing conditions were 37°C and 5% CO2 with saturating humidity – control experiments were performed in parallel to MPO treatments with the cells cultured to the same cells density. MPO treatment was performed with protein purified from human blood (Planta) and at 1 μg/ml final concentration. MPO was inactivated as previously described by Paumann-Page and colleagues (Paumann-Page *et al*, 2013). In brief, 100 μg of MPO was incubated in 0.5 M H2O2 30 min at 37°C, remaining H2O2 was removed by dialysis. Loss of enzymatic activity was confirmed by 3’-(p-aminophenyl) Fluorescein (APF) assay (ThermoFisher).

### Assay for transposase-accessible chromatin using sequencing (ATAC-seq) and data analysis

ATAC-Sequencing was performed with help of Active Motif ATAC-Seq kit as per manufacturer instructions. HUVEC cells were pre-treated with 1 µg/ml of MPO for 2 or 8 hours; for control experiments media was changed accordingly. Cells were lifted up with Trypsin and counted, 50,000 to 100,000 cells used per experiment. Firstly, cells were pelleted by centrifugation at 500 xg for 5 minutes, at 4 °C. Secondly, cells were washed in ice-cold PBS, and pelleted again. Next, cell pellet was resuspended in ice-cold ATAC Lysis Buffer and transferred into a PCR tube. After another round of centrifugation, cell pellet was resuspended in the Tagmentation Master Mix. The tagmentation reaction was performed at 37°C for 1 hour. Further, tagmented DNA was purified. Library preparation was performed as suggested by kit manufacturer. Ready libraries were purified with SPRI clean-up beads. Sequencing was performed in paired-end way and 100 bp per read. Reads were mapped to human genome version hg38. Peak calling was performed with help of MACS2 and Genrich (https://github.com/jsh58/Genrich), differential chromatin accessibility was assessed by csaw R package as described by Reske et al. (Reske *et al*, 2020). To estimate nucleosome positioning around the MPO peaks, reads were sub grouped by length using deepTools package (Ramírez *et al*, 2016) and bamcoverage option with length windows of 10 to 50 bp for short free DNA, 51 to 100 for long free DNA and 180 to 247 for mononucleosomes, exact scaling and RPKM normalization options on.

### Generation and analyses of total RNA-seq

Cells at 2 different time points after treatment with MPO were harvested in Trizol (Life Technologies) and total RNA was isolated and DNase-treated using the Direct-zol RNA miniprep kit (Zymo Research) as per the manufacturer’s instructions. Barcoded cDNA libraries were generated using the TruSeq RNA library kit (Illumina) via selection on poly(dT) beads. The resulting libraries were paired-end sequenced to >50 million read pairs on a HiSeq4000 platform (Illumina). Reads were trimmed and mapped to human genome hg19 using STAR; Reads uniquely mapping to exons were quantified using HTSeq-count and differential gene expression was assessed using DESeq2 (Love *et al*, 2014). Differentially-regulated genes per each cell line and treatment are listed in Table S3.

### MPO immunoprecipitation and proteomics

Immunoprecipitation was performed from isolated nuclei (after 8 hours MPO treatment at a concentration 1 µg/ml), non-treated cells were used as negative control. For immunoprecipitation antibody against MPO (Calbiochem, 475915) was used for both treated and untreated cells. Cell nuclei were isolated by incubating cells for 15 min on ice in NIB buffer (15 mM Tris-HCL pH 7.5, 60 mM KCl, 15 mM NaCl, 5 mM MgCl2, 1 mM CaCl2, 250 mM sucrose) containing 0.3% NP-40 as previously described in (Kargapolova *et al*, 2021b). Nuclei were pelleted for 5 min 800 ×g at 4 °C, washed twice in the same buffer, lysed for 10 min on ice in IP buffer (150 mM LiCl, 50 mM Tris-HCl pH 7.5, 1 mM EDTA, 0.5% Empigen) freshly supplemented with 2 mM sodium vanadate, 1× protease inhibitor cocktail (Roche), PMSF (10 µl), 0,5 mM DTT, and 50 units Benzonase per ml of IP buffer, before preclearing cell debris by centrifugation at >15,000 ×g at 4 °C. Finally, 1 mg of the lysate was incubated with anti-MPO antisera overnight at 4 °C. Magnetic beads (Active Motif) were then washed once with 1× PBS-Tween and combined with the antibody-lysate mixture. Following a 2h incubation at 4 °C, beads were separated on a magnetic rack and washed 5×, 5 min each in wash buffer (150 mM KCl, 5 mM MgCl2, 50 mM Tris-HCl pH 7.5, 0.5% NP-40) and another two times in wash buffer without NP-40. Captured proteins were pre-digested and eluted from the beads using digestion buffer (2 M Urea, 50 mM Tris-HCl pH 7.5, 1 mM DTT) supplemented with trypsin and eluted from the beads with elution buffer (2 M Urea, 50 mM Tris-HCl pH 7.5, 5 mM chloroacetamide) supplemented with trypsin and LysC, before subjected to mass-spectrometry on a Q-Exactive Plus Orbitrap platform coupled to an EASY nLC (Thermo Scientific). Peptides were loaded in solvent A (0.1% formic acid in water) onto an in-house packed analytical column (50 cm length, 75 µm I.D., filled with 2.7 µm Poroshell EC120 C1; Agilent); were chromatographically separated at a constant flow rate of 250 nL/min using the following gradient: 3–8% solvent B (0.1% formic acid in 80% acetonitrile) for 1 min, 8–30% solvent B for 39 min, 30–50% solvent B for 8 min, 50–95% solvent B for 0.3 min, followed by washing and column equilibration. The mass spectrometer was operated in data-dependent acquisition mode. An MS1 survey scan was acquired from 300–1750 m/z at a resolution of 70,000. The top 10 most abundant peptides were isolated within a 1.8 Th window and subjected to HCD fragmentation at a normalized collision energy of 27%. The AGC target was set to 5e5 charges, allowing a maximum injection time of 110 ms. Product ions were detected at a resolution of 35,000; Precursors were dynamically excluded for 10 s. All raw data were processed with Maxquant (v1.5.3.8) using default parameters. Briefly, MS2 spectra were searched against the Uniprot HUMAN fasta (16.06.2017) database, including a list of common contaminants. False discovery rates on were estimated by a target-decoy approach to 1% (Protein FDR) and 1% (PSM FDR), respectively. The minimal peptide length was set to 7 amino acids and carbamidomethylation at cysteine residues was considered as a fixed modification. Oxidation (M) and Acetyl (Protein N-term) were included as variable modifications. The match-between runs option was enabled and LFQ quantification was enabled under default settings. The full list of peptide hits and their analysis is provided in Table S4.

### Immunostaining and imaging

Cells were grown on coverslips, fixed in 4% PFA/PBS for 10 min at room temperature, washed in 1× PBS, permeabilized in 0.5% Triton-X/PBS for 5 min at room temperature, blocked in 1% BSA/PBS for 1 h before incubating with the primary antibody of choice for 2 h to overnight. Cells were next washed twice in 1x PBS for 5 min, before incubating with the appropriate secondary antisera for 1 h at room temperature. Nuclei were counterstained with DAPI (Sigma-Aldrich) for 5 min, washed, and coverslips mounted onto slides in Prolong Gold Antifade (Invitrogen). For image acquisition, a fluorescent microscope Olympus IX83, 60× (Oil) objective was used, making sure exposure times were maintained constant across samples in each imaging session for the same immunostaining. Finally, images were analyzed using the Fiji suite (Schindelin *et al*, 2012) as follows: first, background signal levels were subtracted using the embedded function (rolling ball function of 50-px radius with a sliding paraboloid and disabled smoothing), and the DAPI channel was used to determine the area of interest where signal would be quantified from. Measured mean signal intensities were used to generate plots in R or via InstantClue (Nolte *et al*, 2018). For bean plots, dots represent the mean of the dataset; for box plots, whiskers ends represent the top and bottom quantiles, respectively.

### Nuclei and chromatin fractionation and Western blotting

For nuclei fractionation HUVECs from 15 cm plates were washed in 10 ml/ plate ice-cold PBS and collected by centrifugation (3000g, 3 min, 4 °C). Collected cells were washed with PBS, pelleted and lysed in 1 ml ChIP prep buffer ChIP-IT High Sensitivity kit, Active Motif), supplemented with protease inhibitor cocktail and PMSF. Lysis was performed 10 minutes on ice, following mild sonication as described in NEXSON protocol for nuclei isolation (Arrigoni *et al*, 2015). Nuclei were collected via centrifugation for 3 min at 1250 g at 4 °C and x1 with ChIP prep buffer.

For chromatin fractionation HUVECs were grown to 80-90% confluency, washed with PBS and Trypsinized. Trypsin was inactivated by adding 10% FBS containing media. Cells from x 10 cm dishes were pelleted and lysed in 300 μl of chromatin extraction buffer (20mM Tris, pH 7.5, 100mM NaCl, 5 mM MgCl2, 10% glycerol, 0.2% NP-40, 1mM NaF, 0.5 mM CTT, 1x complete protease inhibitor and phosphatase inhibitor (Roche). Lysate was sonicated using Active Motif dounce homogenizer, with 3 pulses at 25% intensity 30 sec on and 30 sec off. Cell debris was pelleted for 5 minutes at 2500 rpm at 4°C, supernatant was collected. Supernatant from previous step was centrifuged 10 minutes at 13,000g at 4 °C, pellet was washed 3 times using chromatin extraction buffer, this fraction constitutes the chromatin fraction. The supernatant was collected and constitutes soluble fraction. Chromatin fraction was resuspended in 50 μl TBS/Tween with addition 5 μl of benzonase (500 kU/μl, Merk Millipore) and incubated at 37°C 30 minutes, following western blot analysis. Western blotting was performed as previously described (Zirkel *et al*, 2018). In brief, ∼2 × 10^6^ cells were enzymatically detached or gently scraped off cell culture dishes, and pelleted for 5 min at 600 ×g. The supernatant was discarded, and the pellet resuspended in 100 µl of ice-cold RIPA lysis buffer containing 1× protease inhibitor cocktail (Roche), incubated for 30 min on ice, and centrifuged for 15 min at >15,000 ×g to pellet cell debris and collect supernatant. Total protein concentrations were determined using the Pierce BCA Protein Assay Kit (ThermoFisher Scientific), before extracts were stored at −80 °C. Proteins were resolved by SDS-PAGE, transferred onto membranes using the TransBlot Turbo setup (Bio-Rad), and detected using the antibodies and dilutions listed in Table S6.

### MNase treatment and gel electrophoresis, *in vitro* migration assays

HUVECs were grown in 6-well plates to ∼80% confluency and rinsed with PBS prior to addition of 1 ml of freshly prepared permeabilization buffer (15 mM Tris/HCl pH 7.6; 60 mM KCl; 15 mM NaCl; 4 mM CaCl2; 0.5 mM EGTA; 300 mM sucrose; 0.2% NP-40; 0.5 mM β-mercaptoethanol supplemented with MNase (Sigma) at 1 μ/ml final concentration. MNase was added for 2, 4 and 6 min at 37 °C, and stopped by addition of an equal volume of stop buffer (50 mM Tri/HCl pH 8.0; 20 mM EDTA; 1% SDS). Finally, 250 µg RNase A were added for 2 h at 37 °C, followed by addition of 250 µg proteinase K and incubation at 37 °C overnight. Next day, DNA was isolated via standard phenol-chloroform extraction, digestion efficiency was determined after electrophoresis in 1% agarose gels. Images were quantified using Fiji suite.

For *in vitro* migration (scratch) assays, HUVECs were grown in 6-well plates and one scratch per well was manually inflicted using a sterile cell scraper. Cell migration into the scratch was monitored for up to 20 h by bright field microscopy.

### Electromobility shift assay

Nucleosomes were reconstituted as previously described by Luger et al. (Luger *et al*, 1999) and Dyer et al. (Dyer *et al*, 2004). Binding of reconstituted nucleosomes with MPO was performed 30-45 minutes in total volume of 10 μl of binding buffer (25mM HEPES, 150 mM KCl) at room temperature. Titers of MPO varying from 50 nM to 1000 nM were used to bind with 100 nM nucleosomes with short and long linker DNA. Sucrose solution at a final concentration of 5% was added as a loading buffer prior to sample loading. 5% TBE native polyacrylamide gel was used for analysis. Gel was pre-run without samples at 150 V for 60 minutes in the cold room on ice, the run with samples was performed with same voltage settings, for 50 minutes. Gel was stained in 50 ml TBE with 1:10,000 diluted SYBR gold for exactly 10 minutes, washed with water and visualized using GelDoc imaging System (Bio-Rad).

### siRNA experiments ILF3 KD, 3’end sequencing, transcription inhibition by DRB

Universal negative control siRNA (Mission siRNA Universal Negative Control #1, SIC001, Sigma) and ILF3 siRNA1 and ILF3siRNA2 (ILF3_1 PDSIRNA2D, SASI_Hs01_00242960, SASI_Hs01_00242961, Sigma) were used in ILF3 knockdown experiments. RNAiMAX (ThermoFisher) and Opti-MEM (ThermoFisher) were used for transfection. To ensure an ideal knockdown rate, HUVECs were cultured in complete Endopan3 medium on 6-well plates (overnight), and shifted to no-antibiotic Endopan3 medium for siRNA transfection. Transfection was done according to the protocol provided by ThermoFisher. Knockdown efficiency was assessed by qPCR 24 hours and by western blot 72 hours after transfection. 3’-end RNA-seq was used as a lower-cost alternative to mRNA-seq. Libraries were prepared from total RNA using the QuantSeq 3’ mRNA-Seq Library Prep Kit (Lexogen), and paired-end sequenced on a HiSeq4000 platform (Illumina) generating ∼15 × 10^6^ 100 nt-long reads per sample. Reads were quality assessed and mapped to hg19 using STAR (Dobin *et al*, 2013). Reads uniquely mapping to exons were quantified using HTSeq-count and differential gene expression was assessed using DESeq2 (Love *et al*, 2014). Differentially-regulated genes per each cell line and treatment are listed in Table S5. DRB treatment was performed in triplicates, for 30 min or 1 hour on ECs, cultured on 6- well plates. 3.1 mM DRB (TCI chemicals) stock solution in DMSO was applied to ECs at final concentration of 0.1 mM. Prior to DRB treatment cells were treated with siRNA against ILF3 or control siRNA for 71 hour, MPO treatment was performed for 2 hours at a standard concentration of 1 µg/ml. Cells were collected at 72 hours post-transfection, RNA was isolated, subjected to cDNA synthesis and qPCR.

### Left Anterior Descending Artery Ligation and DAB staining

LAD ligation was performed as previously described (Guthoff *et al*, 2022). Mice were anesthetized with isoflurane, received low dose buprenorphine subcutaneously (Essex-Pharma, Munich, Germany; 0.05 mg/kg bodyweight) for analgesia and were placed on a heating pad to regulate body temperature. Following endotracheal intubation, animals were ventilated with 150 strokes/min and stroke volume of 7 μl/g bodyweight (Harvard Apparatus, Holliston, MA, USA). Surgical procedures were carried out using a dissecting microscope (Leica MZ6, Leica Microsystems, Wetzlar, Germany). After lateral thoracotomy of the fourth intercostal space, a suture (8/0 polypropylene suture, Polypro, CP Medical, Norcross, GA, USA) was placed around the LAD and the artery was ligated with a bow tie. Ischemia was visually confirmed by blanching of the left ventricular (LV) apex. Mice were sacrificed 8 weeks after infarction, hearts were processed for DAB staining using DAB staining kit (Abcam, ab64238) and following the manufacturer recommendations, at least 50 images per mouse were analysed using Qupath software (Bankhead *et al*, 2017).

### Animal Studies and Ethics Statement

Male, 8- to 12-week-old Mpo^-/-^ and wild type (WT) mice were used for MI. Animals were of C57BL/6J background (Jackson Laboratory, Bar Harbor, ME, USA). The strategy for the generation of Mpo ^-/-^ mice has been described previously (Brennan *et al*, 2001). All animal studies were approved by the local authorities (State Agency for Nature, Environment and Consumer Protection (LANUV), Recklinghausen, NRW, Germany) and the University Cologne Animal Care and Use Committees. All surgical interventions were performed under anaesthesia using isoflurane and perioperative analgesia with buprenorphine to minimize suffering.

### Statistics and reproducibility

P-values associated with Student’s/Welch t-tests were calculated using GraphPad [http://graphpad.com/], and those associated with the Wilcoxon-Mann-Whitney test using the EDISON-WMW [https://ccb-compute2.cs.uni-saarland.de/wtest/] online tool. Unless otherwise stated, only P-values <0.05 were deemed as significant. Also, note that for immunofluorescence analyses, representative images from one of at least two independent and converging experiments are displayed and quantified.

## ACCESSION NUMBERS

Proteomics data are available via ProteomeXchange with identifier PXD033079. Sequencing data are available via GEO with identifier GSE202015.

The following ENCODE data were analysed:

POLR2A ChIP (https://www.encodeproject.org/experiments/ENCSR000EFB/)

H3K27me3 ChIP (https://www.encodeproject.org/experiments/ENCSR000DVO/)

H3K4me3 ChIP (https://www.encodeproject.org/experiments/ENCSR000DVN/)

Control (input) for ChIP (https://www.encodeproject.org/experiments/ENCSR000EVV/)

H3K4me3 ChIP (https://www.encodeproject.org/experiments/ENCSR000AKN/)

H3K4me1 ChIP (https://www.ncbi.nlm.nih.gov/geo/query/acc.cgi?acc=GSM733690)

H3K27ac ChIP (https://www.ncbi.nlm.nih.gov/geo/query/acc.cgi?acc=GSM733691)

H3K27ac ChIP (https://www.ncbi.nlm.nih.gov/geo/query/acc.cgi?acc=GSM3557982)

H3K4me3 ChIP (https://www.encodeproject.org/experiments/ENCSR000AKN/)

## SUPPLEMENTARY DATA

Table S1 - MPO and iMPO ChIP-Seq, overlap with regulated gene set

Table S2 - ATAC-seq regions of increased chromatin / decreased chromatin accessibility

Table S3 - RNA-Seq 2h MPO, 8h MPO, Q91T, TNF-alpha

Table S4 - MPO Co-IP MS data, protein phosphorylation analysis

Table S5 - 3’-end sequencing data

Table S6 – List of Antibodies and Primers used in this study

## ACKNOWLEDGEMENTS

We would like to thank all members of the Papantonis, Rada-Iglesias, Poepsel and Baldus laboratories for helpful discussions. We also thank the CMMC Imaging and Embedding facilities, the CECAD Proteomics facility, the Cologne Center for Genomics, the High-Performance Computing cluster “CHEOPS” of the University of Cologne for continuous access to resources. We thank Prof. Paul Georg Furtmüller for supplying us with the Q91T mutant MPO protein.

## FUNDING

This work was supported by grants from the German Research Foundation to S.B. and A.P. (CCRC 2407– 360043781) to S.B and M.A. (CRC TRR259–397484323), Y.K. was supported by the Koeln Fortune Program, Faculty of Medicine, University of Cologne.

## AUTHOR CONTRIBUTIONS

Y.K., A.P., S.P., S.B. conceived the study. R.Z., K.M., E.P., S.G., S.G., H.M., R.R. performed experiments. T.G., K.M., Y.K. performed bioinformatic analysis. R.Z., R.R., J.B. performed and analyzed imaging experiments. J.-W.L. analysed mass-spectrometry data. J.A., P.N. contributed with NGS library preparation and sequencing. A.R.-I., S.B., A.P., M.A. were responsible for funding acquisition. Y.K., A.P., R.Z., S.P., S.B. drafted the manuscript with contributions of all authors.

## CONFLICT OF INTEREST

The authors declare no conflict of interest.

## SUPPLEMENTARY FIGURES

**Figure S1.**
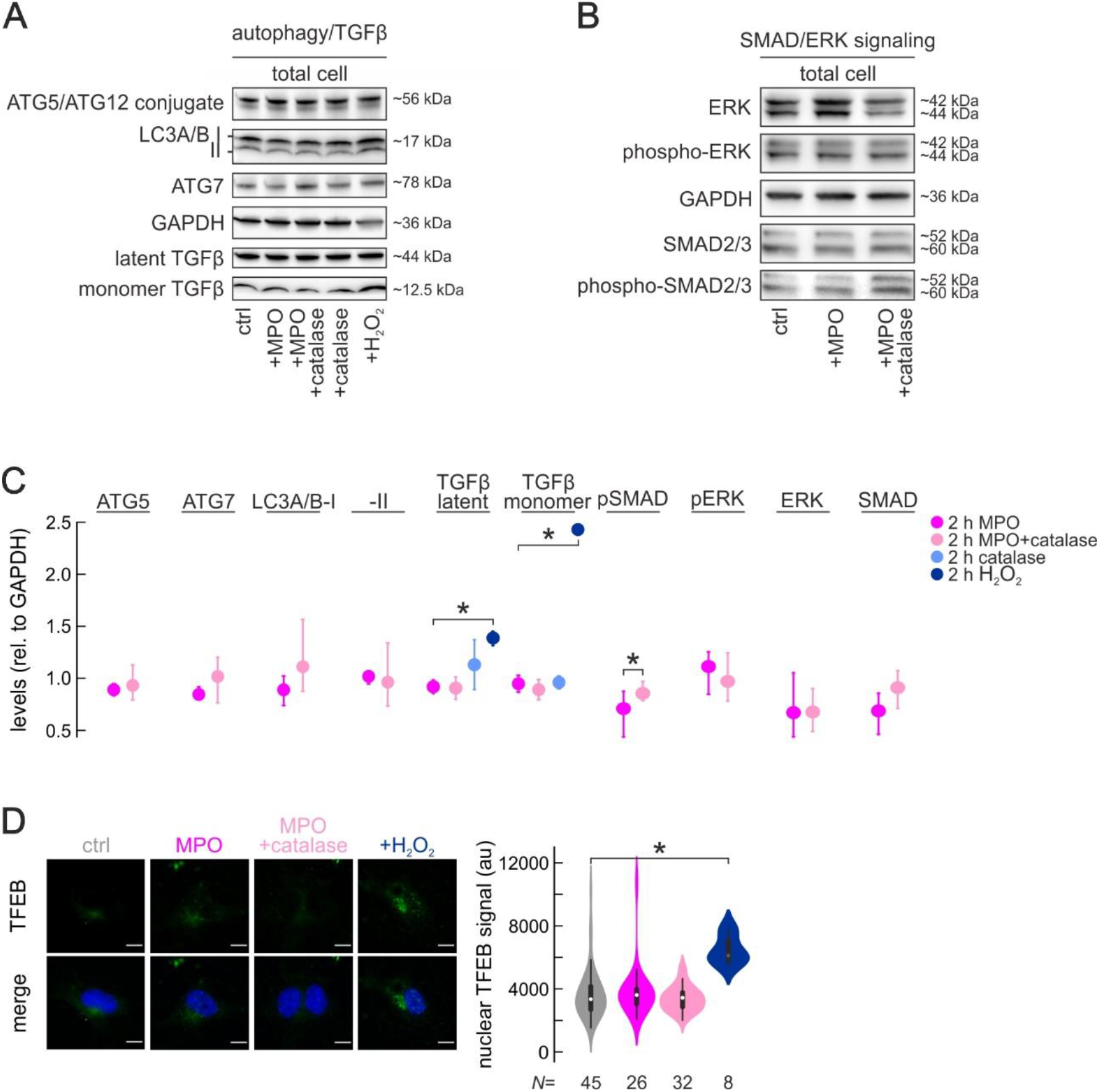
MPO translocates into the cytoplasm and nuclei of endothelial cells. (A) Western blot represents levels of ATG5-ATG12, LC3A/B, ATG7, GAPDH, latent TGFß and monomer TGFß upon MPO, MPO and catalase, catalase and H2O2 (used as positive control) treatments in ECs, representative images of three biological replicates. (B) Western blot represents levels of SMAD2/3, phospho-SMAD2/3, ERK, phospho-ERK, GAPDH upon 2 hour MPO treatment or combination of MPO and catalase of ECs, representative image of three replicates. (C) Relative levels of protein depicted on Figure S1A and S1B, normalized to GAPDH, data are presented as mean ±SEM. (D) Representative TFEB immunofluorescence images of ECs after 2 hours of MPO treatment, combination of MPO with catalase treatment or H2O2 (used as positive control to confirm TFEB translocation as shown by Xudong Sun et al. 2021) and control media (scale bar=10μm). Quantification of IF images (from 2 independent experiments) are depicted with help of violin plot (right panel).

**Figure S2.**
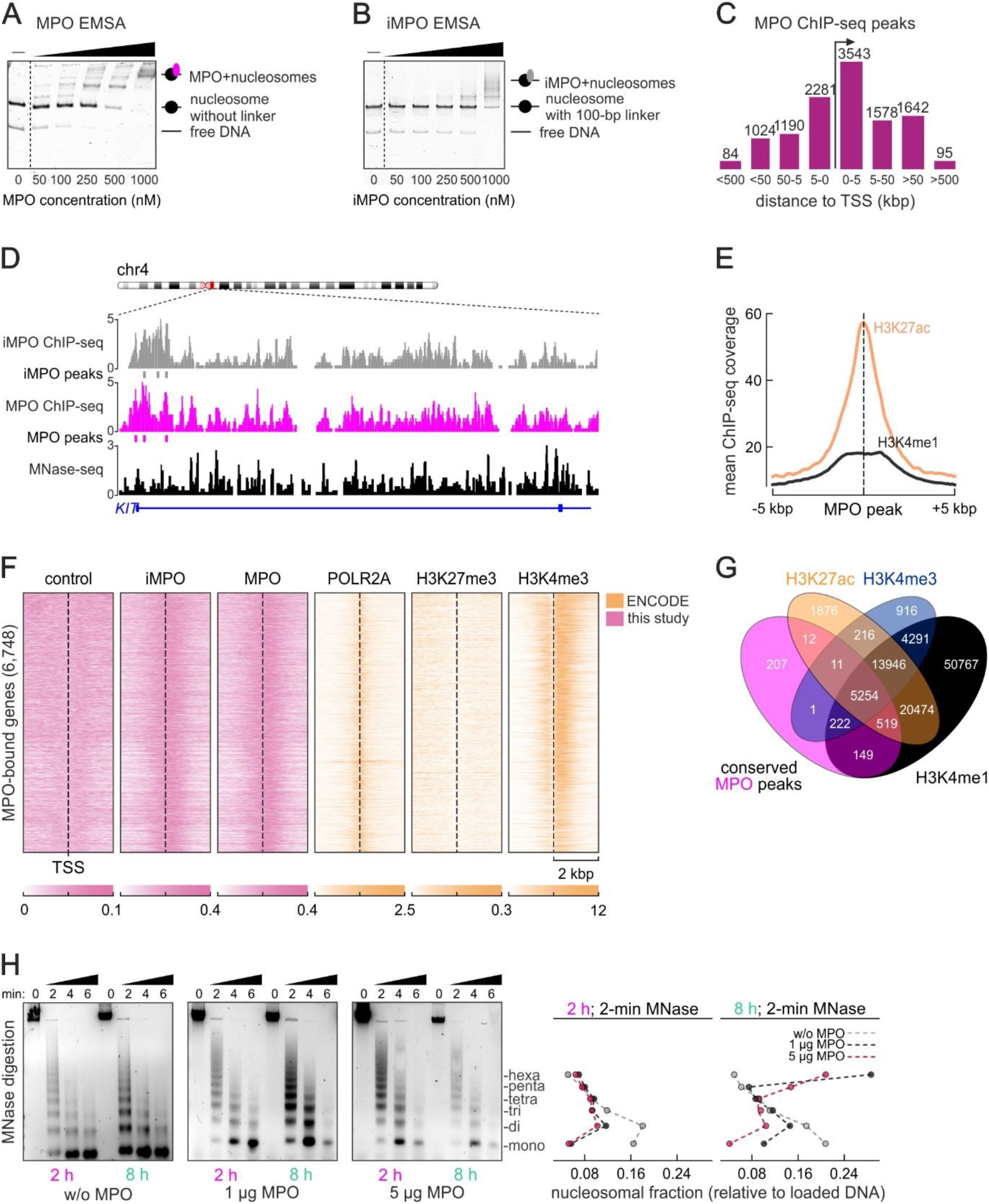
Binding of Myeloperoxidase to chromatin, sites distribution, IDR analysis and chromatin condensation. (A) DNA EMSAs performed with increasing titers of MPO with ‘no linker’ DNA nucleosomes. (B) DNA EMSAs performed with increasing titers of inactivated iMPO and long linker DNA nucleosomes (100bp). (C) Barplots showing distribution of MPO binding sites around transcription start sites (TSS) of bound genes. (D) Representative genome browser view of MPO and iMPO ChIP-Seq in MPO/iMPO-treated HUVEC cells (upper panels, current work) and MNase-Seq experiment by Diermeier and colleagues (Diermeier *et al*, 2014). (E) H3K27ac and H3K4me1 ChIP-Seq (ENCODE) coverage centered at around MPO peaks shows high levels of H3K27ac signal. (F) Heatmaps showing input, iMPO, MPO (current study), POLR2A, H3K27me3, H3K4me3 (ENCODE) ChIP-Seq signal in the 4 kbp windows around MPO-bound TSS. (G) Venn diagram depicts an overlap between conserved MPO peaks (reproducible between two biological replicates), H3K27ac, H3K4me3 and H3K4me1 marks. (H) Agarose gel images resolve MNase-treated DNA fragments, corresponding to mono-, di-, tri-, tetra, penta- and hexa-nucleosomes after 2 and 8 h MPO treatment or mock treatment. The abundance of each sort of fragments is quantified and normalized to the total DNA loaded onto the gel, and plotted as a line plot.

**Figure S3.**
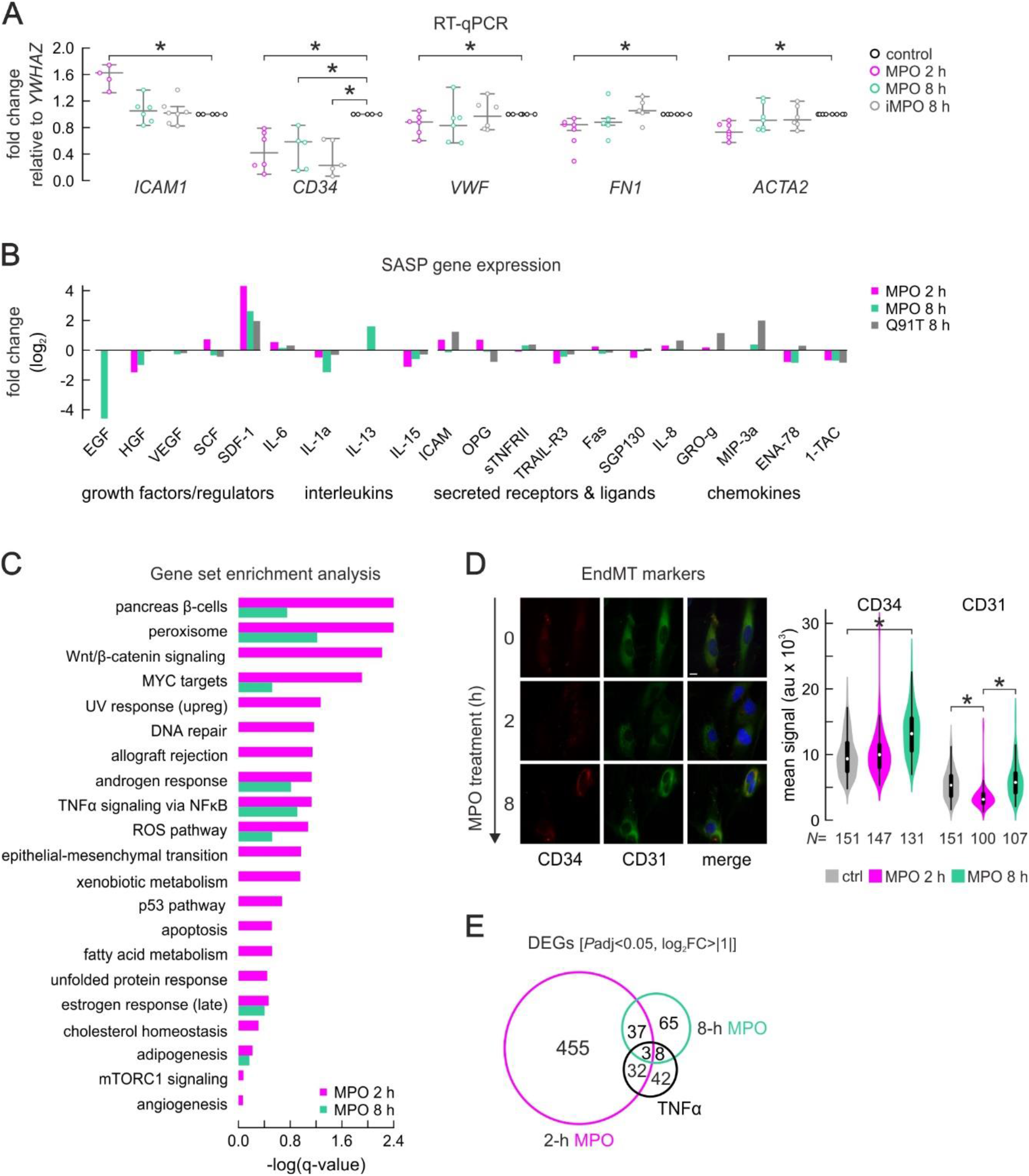
Signalling pathways regulated after 2 and 8 h of MPO treatment. (A) Relative expression of *Icam1*, *CD34 Vwf*, *Fn1* and *Acta2*, upon MPO (2h and 8h) and iMPO (8h) treatment, measured by qPCR and normalized to housekeeping transcript *Ywhaz* and relative to non-treated cells. Data are presented as mean ±SEM, asterisk labels Welch-test signif. pval<0.05. (B) Gene expression changes (RNA-Seq) of a subset of genes, grouped as growth factors, interleukins, secreted ligands, chemokines, represented as bars, showing log2 fold changes between wild type MPO (2 and 8 h) or Q91T mutant MPO (8h) treated samples and controls. (C) Gene set enrichment analysis (GSEA) of ranked gene expression data, generated by RNA-Seq of HUVECS under 2 and 8 hour MPO treatment and control (fresh media) treatment. Bar plots represent –log10 FDR q-values of GSEA results for 2 (pink) and 8 (green) hour MPO-treated samples. (D) Representative immunofluorescence images of markers of Endothelial-to-Mesenchymal transition CD31and CD34 in ECs after 0, 2 and 8 hours of treatment, scale bar=10μm. Treatments performed in triplicates and violin plots depict quantifications of 10-15 random images selected for each biological replica. (E) Venn diagrams showing an overlap between differentially expressed genes after 2 and 8 hours MPO and TNFα treatment (Diermeier *et al*, 2014).

**Figure S4.**
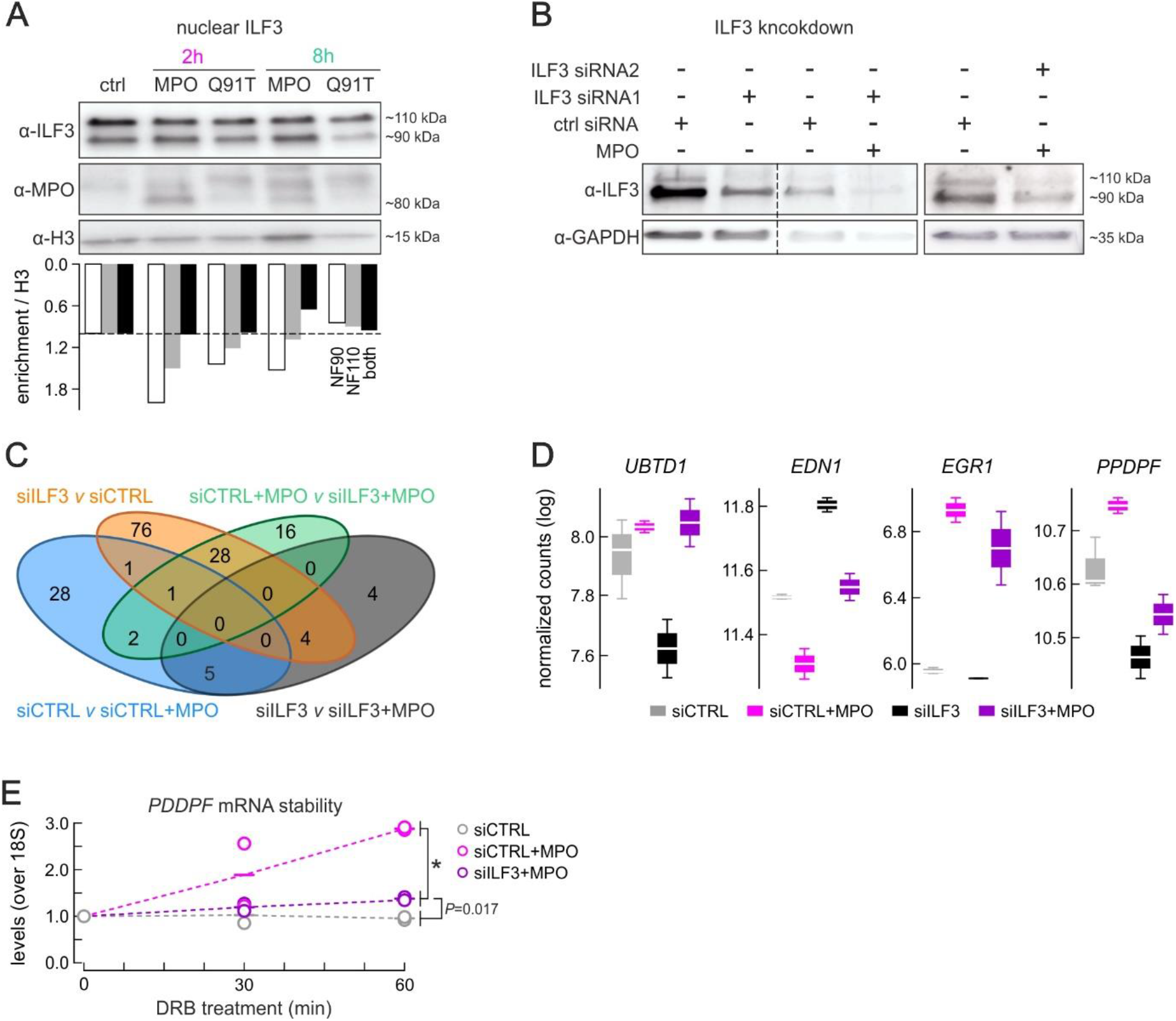
MPO treatment triggers changes in ILF3/NF90 levels in cytoplasm and nucleus and empowers regulation of ILF3-bound mRNAs. (A) Western blot image performed on nuclear fraction of ECs upon treatment with MPO or Q91T mutant MPO variant, and showing ILF3/NF90 increment in the nucleus post- treatment. Quantification of Western blot, normalized to histone H3 levels (lower panel). (B) Western blot image, controlling for high efficacy of ILF3 depletion, performed with 2 different siRNAs against ILF3 and accompanied by MPO treatment prior to 3’-end Sequencing experiment. (C) Venn diagram shows an overlap between 4 gene sets (generated by 3’-end RNA-sequencing) including differentially expressed upon MPO treatment and with or without ILF3 depletion. (D) Box plots show mean regularized log transformed counts (generated by 3’-end RNA-sequencing) for a subset genes, whose expression is regulated by ILF3 depletion and restored by co-treatment with MPO (*UBTD1*, *EDN1*), and a subset of genes regulated by MPO independently of ILF3 (*EGR1*) or restored by combination of MPO treatment with ILF3 depletion (*PPDPF*).

**Figure S5.**
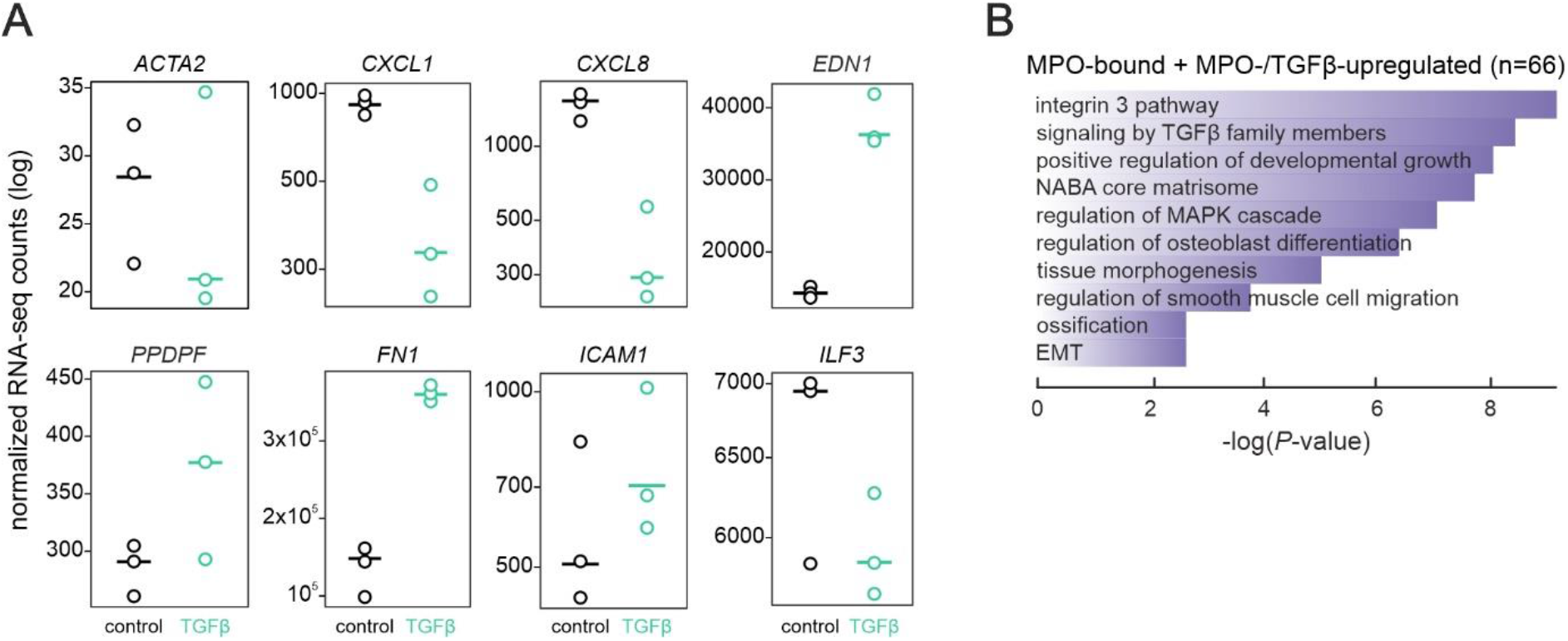
Overlapping gene regulation by MPO and TGFβ are bound by MPO and involved in EMT. (A) Dotplot, showing differential expression of selected transcripts after TGFβ treatment in HUVECs. (B) Metascape analysis of genes bound by MPO (+/- 1000 bp from the peak center) and regulated by either MPO or TGFβ or both.

## REFERENCES

Abhinand CS, Raju R, Soumya SJ, Arya PS & Sudhakaran PR (2016) VEGF-A/VEGFR2 signaling network in endothelial cells relevant to angiogenesis. J Cell Commun Signal 10: 347–354

Arrigoni L, Richter AS, Betancourt E, Bruder K, Diehl S, Manke T & Bonisch U (2015) Standardizing chromatin research: A simple and universal method for ChIP-seq. Nucleic Acids Res 44

Astern JM, Pendergraft WF, Falk RJ, Jennette JC, Schmaier AH, Mahdi F & Preston GA (2007) Myeloperoxidase interacts with endothelial cell-surface cytokeratin 1 and modulates bradykinin production by the plasma kallikrein-kinin system. American Journal of Pathology 171: 349–360

Baldus S, Eiserich JP, Mani A, Castro L, Figueroa M, Chumley P, Wenxin M, Tousson A, Roger White C, Bullard DC, et al (2001) Endothelial transcytosis of myeloperoxidase confers specificity to vascular ECM proteins as targets of tyrosine nitration. Journal of Clinical Investigation 108: 1759–1770

Bankhead P, Loughrey MB, Fernández JA, Dombrowski Y, McArt DG, Dunne PD, McQuaid S, Gray RT, Murray LJ, Coleman HG, et al (2017) QuPath: Open source software for digital pathology image analysis. Sci Rep 7: 16878

Bischoff J, Casanovas G, Wylie-Sears J, Kim DH, Bartko PE, Guerrero JL, Dal-Bianco JP, Beaudoin J, Garcia ML, Sullivan SM, et al (2016) CD45 Expression in Mitral Valve Endothelial Cells after Myocardial Infarction. Circ Res 119: 1215–1225

Bradley PP, Christensen RD & Rothstein G (1982) Cellular and extracellular myeloperoxidase in pyogenic inflammation. Blood 60: 618–622

Brennan ML, Anderson MM, Shih DM, Qu XD, Wang X, Mehta AC, Lim LL, Shi W, Hazen SL, Jacob JS, et al (2001) Increased atherosclerosis in myeloperoxidase-deficient mice. Journal of Clinical Investigation 107: 419– 430

Castella S, Bernard R, Corno M, Fradin A & Larcher JC (2015) Ilf3 and NF90 functions in RNA biology. Wiley Interdiscip Rev RNA 6: 243–256

Davies MJ (2011) Myeloperoxidase-derived oxidation: mechanisms of biological damage and its prevention. J Clin Biochem Nutr 48: 8

Davies MJ & Hawkins CL (2020) The Role of Myeloperoxidase in Biomolecule Modification, Chronic Inflammation, and Disease Antioxid Redox Signal

Diermeier S, Kolovos P, Heizinger L, Schwartz U, Georgomanolis T, Zirkel A, Wedemann G, Grosveld F, Knoch TA, Merkl R, et al (2014) TNFα signalling primes chromatin for NF-κB binding and induces rapid and widespread nucleosome repositioning. Genome Biol 15: 536

Dobin A, Davis CA, Schlesinger F, Drenkow J, Zaleski C, Jha S, Batut P, Chaisson M & Gingeras TR (2013) STAR: Ultrafast universal RNA-seq aligner. Bioinformatics 29: 15–21

Dyer PN, Edayathumangalam RS, White CL, Bao Y, Chakravarthy S, Muthurajan UM & Luger K (2004) Reconstitution of nucleosome core particles from recombinant histones and DNA. Methods Enzymol 375: 23–44

Edwards SW, Hughes V, Barlow J & Bucknall R (1988) Immunological detection of myeloperoxidase in synovial fluid from patients with rheumatoid arthritis. Biochemical Journal 250: 81–85

Glaser SF, Heumüller AW, Tombor L, Hofmann P, Muhly-Reinholz M, Fischer A, Günther S, Kokot KE, Hassel D, Kumar S, et al (2020) The histone demethylase JMJD2B regulates endothelial-to-mesenchymal transition. Proc Natl Acad Sci U S A 117: 4180–4187

Guthoff H, Hof A, Klinke A, Maaß M, Konradi J, Mehrkens D, Geißen S, Nettersheim FS, Braumann S, Michaelsson E, et al (2022) Protective Effects of Therapeutic Neutrophil Depletion and Myeloperoxidase Inhibition on Left Ventricular Function and Remodeling in Myocardial Infarction. Antioxidants (Basel) 12

Kargapolova Y, Geißen S, Zheng R, Baldus S, Winkels H & Adam M (2021a) The enzymatic and non-enzymatic function of myeloperoxidase (Mpo) in inflammatory communication. Antioxidants 10 doi:10.3390/antiox10040562 [PREPRINT]

Kargapolova Y, Rehimi R, Kayserili H, Brühl J, Sofiadis K, Zirkel A, Palikyras S, Mizi A, Li Y, Yigit G, et al (2021b) Overarching control of autophagy and DNA damage response by CHD6 revealed by modeling a rare human pathology. Nat Commun 12: 3014

El Kebir D, József L, Pan W & Filep JG (2008) Myeloperoxidase delays neutrophil apoptosis through CD11b/CD18 integrins and prolongs inflammation. Circ Res 103: 352–359

Khalil A, Medfai H, Poelvoorde P, Kazan MF, Delporte C, Van Antwerpen P, EL-Makhour Y, Biston P, Delrée P, Badran B, et al (2018) Myeloperoxidase promotes tube formation, triggers ERK1/2 and Akt pathways and is expressed endogenously in endothelial cells. Arch Biochem Biophys 654: 55–69

Kovacic JC, Dimmeler S, Harvey RP, Finkel T, Aikawa E, Krenning G & Baker AH (2019) Endothelial to Mesenchymal Transition in Cardiovascular Disease. J Am Coll Cardiol 73: 190

Lefkowitz DL, Roberts E, Grattendick K, Schwab C, Stuart R, Lincoln J, Allen RC, Moguilevsky N, Bollen A & Lefkowitz SS (2000) The endothelium and cytokine secretion: The role of peroxidases as immunoregulators. Cell Immunol 202: 23–30

Liao R & Mizzen CA (2017) Site-specific regulation of histone H1 phosphorylation in pluripotent cell differentiation. Epigenetics Chromatin 10

Liu X, Yu M, Chen Y & Zhang J (2016) Galunisertib (LY2157299), a transforming growth factor-β receptor I kinase inhibitor, attenuates acute pancreatitis in rats. Braz J Med Biol Res 49

Lopes-Paciencia S, Saint-Germain E, Rowell MC, Ruiz AF, Kalegari P & Ferbeyre G (2019) The senescence-associated secretory phenotype and its regulation. Cytokine 117: 15–22

Love MI, Huber W & Anders S (2014) Moderated estimation of fold change and dispersion for RNA-seq data with DESeq2. Genome Biol 15

Luger K, Rechsteiner TJ & Richmond TJ (1999) Preparation of nucleosome core particle from recombinant histones. Methods Enzymol 304: 3–19

Manchanda K, Kolarova H, Kerkenpaß C, Mollenhauer M, Vitecek J, Rudolph V, Kubala L, Baldus S, Adam M & Klinke A (2018) MPO (myeloperoxidase) reduces endothelial glycocalyx thickness dependent on its cationic charge. Arterioscler Thromb Vasc Biol 38: 1859–1867

Metzler KD, Fuchs TA, Nauseef WM, Reumaux D, Roesler J, Schulze I, Wahn V, Papayannopoulos V & Zychlinsky A (2011) Myeloperoxidase is required for neutrophil extracellular trap formation: Implications for innate immunity. Blood 117: 953–959

Moguilevsky N, Garcia-Quintana L, Jacquet A, Tournay C, Fabry L, Piérard L & Bollen A (1991) Structural and biological properties of human recombinant myeloperoxidase produced by Chinese hamster ovary cell lines. Eur J Biochem 197: 605–614

Nolte H, MacVicar TD, Tellkamp F & Krüger M (2018) Instant Clue: A Software Suite for Interactive Data Visualization and Analysis. Sci Rep 8

Nussbaum C, Klinke A, Adam M, Baldus S & Sperandio M (2013) Myeloperoxidase: A leukocyte-derived protagonist of inflammation and cardiovascular disease. Antioxid Redox Signal 18: 692–713

Papayannopoulos V, Metzler KD, Hakkim A & Zychlinsky A (2010) Neutrophil elastase and myeloperoxidase regulate the formation of neutrophil extracellular traps. Journal of Cell Biology 191: 677–691

Paumann-Page M, Furtmüller PG, Hofbauer S, Paton LN, Obinger C & Kettle AJ (2013) Inactivation of human myeloperoxidase by hydrogen peroxide. Arch Biochem Biophys 539: 51–62

Piera-Velazquez S & Jimenez SA (2019) Endothelial to mesenchymal transition: Role in physiology and in the pathogenesis of human diseases. Physiol Rev 99: 1281–1324

Pilsczek FH, Salina D, Poon KKH, Fahey C, Yipp BG, Sibley CD, Robbins SM, Green FHY, Surette MG, Sugai M, et al (2010) A Novel Mechanism of Rapid Nuclear Neutrophil Extracellular Trap Formation in Response to Staphylococcus aureus . The Journal of Immunology 185: 7413–7425

Ramírez F, Ryan DP, Grüning B, Bhardwaj V, Kilpert F, Richter AS, Heyne S, Dündar F & Manke T (2016) deepTools2: a next generation web server for deep-sequencing data analysis. Nucleic Acids Res 44: W160– W165

Reichman TW, Parrott AM, Fierro-Monti I, Caron DJ, Kao PN, Lee CG, Li H & Mathews MB (2003) Selective Regulation of Gene Expression by Nuclear Factor 110, a Member of the NF90 Family of Double-stranded RNA-binding Proteins. J Mol Biol 332: 85–98

Reske JJ, Wilson MR & Chandler RL (2020) ATAC-seq normalization method can significantly affect differential accessibility analysis and interpretation. Epigenetics & Chromatin 2020 13:1 13: 1–17

Rudolph TK, Wipper S, Reiter B, Rudolph V, Coym A, Detter C, Lau D, Klinke A, Friedrichs K, Rau T, et al (2012) Myeloperoxidase deficiency preserves vasomotor function in humans. Eur Heart J 33: 1625–1634

Schindelin J, Arganda-Carreras I, Frise E, Kaynig V, Longair M, Pietzsch T, Preibisch S, Rueden C, Saalfeld S, Schmid B, et al (2012) Fiji: An open-source platform for biological-image analysis Nat Methods

Shen L, Shao N, Liu X & Nestler E (2014) Ngs.plot: Quick mining and visualization of next-generation sequencing data by integrating genomic databases. BMC Genomics 15

Teng N, Maghzal GJ, Talib J, Rashid I, Lau AK & Stocker R (2017) The roles of myeloperoxidase in coronary artery disease and its potential implication in plaque rupture. Redox Report 22: 51–73 doi:10.1080/13510002.2016.1256119 [PREPRINT]

Tombor LS, John D, Glaser SF, Luxán G, Forte E, Furtado M, Rosenthal N, Baumgarten N, Schulz MH, Wittig J, et al (2021) Single cell sequencing reveals endothelial plasticity with transient mesenchymal activation after myocardial infarction. Nat Commun 12

Tominaga-Yamanaka K, Abdelmohsen K, Martindale JL, Yang X, Taub DD & Gorospe M (2012) NF90 coordinately represses the senescence-associated secretory phenotype. Aging (Albany NY) 4: 695

Tripathi S, Pohl MO, Zhou Y, Rodriguez-Frandsen A, Wang G, Stein DA, Moulton HM, Dejesus P, Che J, Mulder LCF, et al (2015) Meta- and Orthogonal Integration of Influenza ‘oMICs’ Data Defines a Role for UBR4 in Virus Budding. Cell Host Microbe 18: 723–735

Vrakas CN, Herman AB, Ray M, Kelemen SE, Scalia R & Autieri M V. (2019) RNA stability protein ILF3 mediates cytokine-induced angiogenesis. The FASEB Journal 33: 3304

Wu TH, Shi L, Adrian J, Shi M, Nair R V., Snyder MP & Kao PN (2018) NF90/ILF3 is a transcription factor that promotes proliferation over differentiation by hierarchical regulation in K562 erythroleukemia cells. PLoS One 13: e0193126

Yang JJ, Preston GA, Pendergraft WF, Segelmark M, Heeringa P, Hogan SL, Jennette JC & Falk RJ (2001) Internalization of proteinase 3 is concomitant with endothelial cell apoptosis and internalization of myeloperoxidase with generation of intracellular oxidants. American Journal of Pathology 158: 581–592

Yan H, Evans J, Kalmbach M, Moore R, Middha S, Luban S, Wang L, Bhagwate A, Li Y, Sun Z, et al (2014) HiChIP: A high-throughput pipeline for integrative analysis of ChIP-Seq data. BMC Bioinformatics 15

Yogalingam G, Lee AR, Mackenzie DS, Maures TJ, Rafalko A, Prill H, Berguig GY, Hague C, Christianson T, Bell SM, et al (2017) Cellular uptake and delivery of myeloperoxidase to lysosomes promote lipofuscin degradation and lysosomal stress in retinal cells. Journal of Biological Chemistry 292: 4255–4265

Zhang Y, Liu T, Meyer CA, Eeckhoute J, Johnson DS, Bernstein BE, Nussbaum C, Myers RM, Brown M, Li W, et al (2008) Model-based analysis of ChIP-Seq (MACS). Genome Biol 9

Zirkel A, Nikolic M, Sofiadis K, Mallm JP, Brackley CA, Gothe H, Drechsel O, Becker C, Altmüller J, Josipovic N, et al (2018) HMGB2 Loss upon Senescence Entry Disrupts Genomic Organization and Induces CTCF Clustering across Cell Types. Mol Cell 70: 730–744.

